# Combinatorial transcriptional profiling of mouse and human enteric neurons identifies shared and disparate subtypes *in situ*

**DOI:** 10.1101/2020.07.03.187211

**Authors:** Aaron A. May-Zhang, Eric Tycksen, Austin N. Southard-Smith, Karen K. Deal, Joseph T. Benthal, Dennis P. Buehler, Mike Adam, Alan J. Simmons, James R. Monaghan, Brittany K. Matlock, David K. Flaherty, S. Steven Potter, Ken S. Lau, E. Michelle Southard-Smith

**Affiliations:** Division of Genetic Medicine, Department of Medicine, Vanderbilt University School of Medicine, Nashville, TN, USA; Genome Technology Access Center, McDonnell Genome Institute, St. Louis, MO, USA; Epithelial Biology Center and the Department of Cell & Developmental Biology, Vanderbilt University School of Medicine, Nashville, TN, USA; University of Cincinnati Children’s Medical Hospital Research Center, Cincinnati, OH, USA; Northeastern University, Department of Biology, Boston, MA, USA; Office of Shared Resources, Vanderbilt University School of Medicine, Nashville, TN, USA

**Keywords:** enteric nervous system (ENS), RNA Sequencing (RNA-Seq), single-nucleus RNA-Seq, in situ hybridization chain reaction (HCR)

## Abstract

**BACKGROUND & AIMS:** The enteric nervous system (ENS) coordinates essential intestinal functions through the concerted action of diverse enteric neurons (EN). However, integrated molecular knowledge of EN subtypes is lacking. To compare human and mouse ENs, we transcriptionally profiled healthy ENS from adult humans and mice. We aimed to identify transcripts marking discrete neuron subtypes and visualize conserved EN subtypes for humans and mice in multiple bowel regions.

**METHODS:** Human myenteric ganglia and adjacent smooth muscle were isolated by laser-capture microdissection for RNA-Seq. Ganglia-specific transcriptional profiles were identified by computationally subtracting muscle gene signatures. Nuclei from mouse myenteric neurons were isolated and subjected to single-nucleus RNA-Seq (snRNA-Seq), totaling over four billion reads and 25,208 neurons. Neuronal subtypes were defined using mouse snRNA-Seq data. Comparative informatics between human and mouse datasets identified shared EN subtype markers, which were visualized *in situ* using hybridization chain reaction (HCR).

**RESULTS:** Several EN subtypes in the duodenum, ileum, and colon are conserved between humans and mice based on orthologous gene expression. However, some EN subtype-specific genes from mice are expressed in completely distinct morphologically defined subtypes in humans. In mice, we identified several neuronal subtypes that stably express gene modules across all intestinal segments, with graded, regional expression of one or more marker genes.

**CONCLUSIONS:** Our combined transcriptional profiling of human myenteric ganglia and mouse EN provides a rich foundation for developing novel intestinal therapeutics. There is congruency among some EN subtypes, but we note multiple species differences that should be carefully considered when relating findings from mouse ENS research to human GI studies.

Graphical Abstract

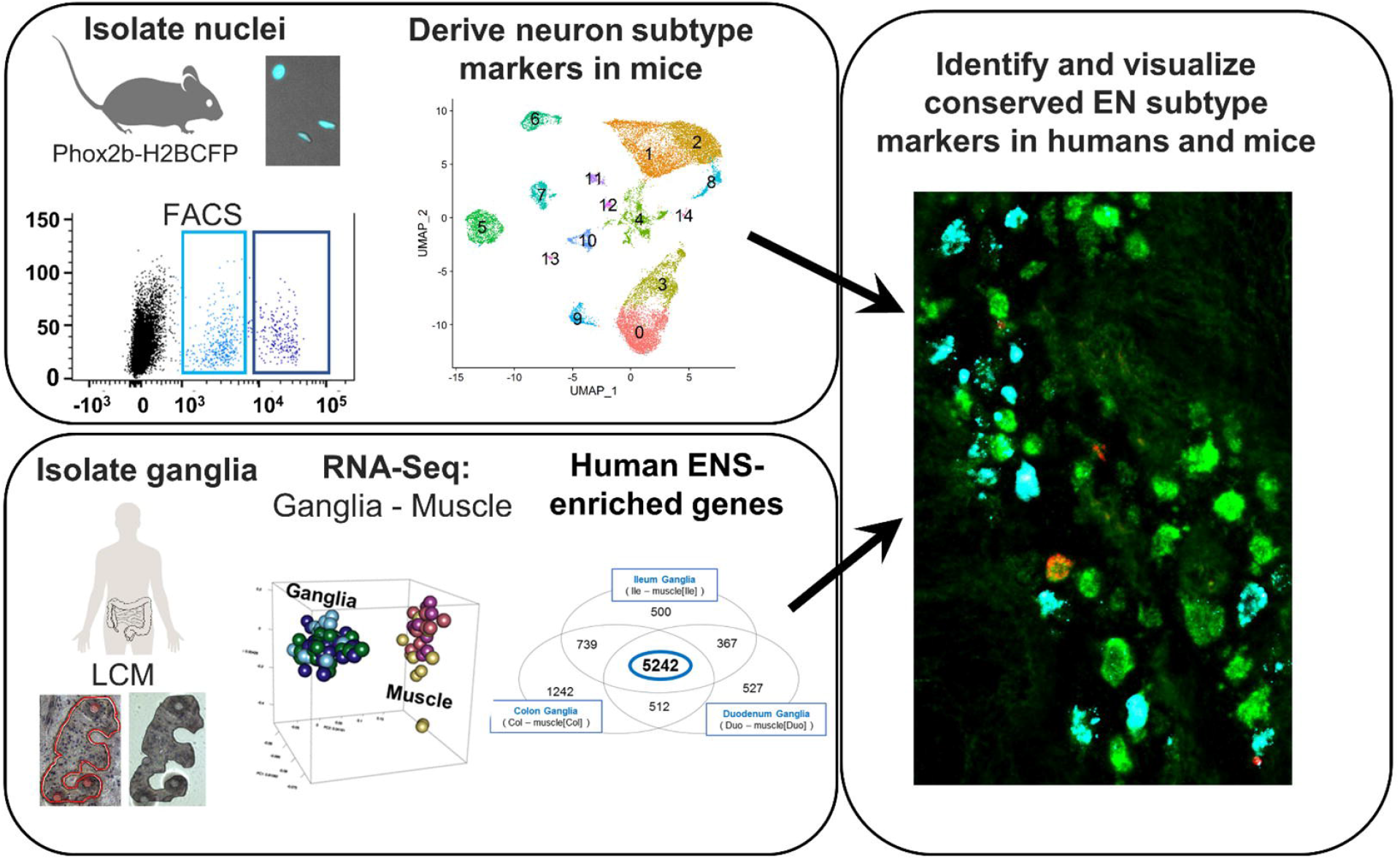

## Introduction

Gastrointestinal motility, osmotic and pH balance, vasodilation, and secretion are all essential functions coordinated by the ENS. These activities are mediated by a diverse array of neurons, outnumbering those in the spinal cord, that are clustered within interconnected ganglia that extend in a continuous network the entire length of the gastrointestinal tract.

Despite a variety of efforts, understanding of human EN diversity is limited. Extensive studies relying on morphological and immunohistochemical characterization have established an initial framework of EN subtypes and identified to date at least nine major neuron classes within human myenteric ganglia ^1, 2^. However, only a handful of immunohistochemical markers label distinct neuron types and even fewer label subtypes reliably across species (i.e. CHAT, NOS1, VIP) ^1^. Because EN morphology is not well-conserved between species, our ability to translate research between humans and rodent models has been constrained. Moreover, prior use of diseased or aged human intestinal tissue has not fulfilled the need for a clinical gold-standard atlas detailing the normal composition and distribution of neuronal subtypes in healthy, young adults.

Rodent models have offered greater access to the inner workings of the ENS and the neuron subtypes mediating its functions. Individual genes that cause discrete deficiencies in ENS development, like Hirschsprung disease or those associated with gastrointestinal motility phenotypes, have been identified by homologous gene knockout or genetic mapping ^3–5^. In addition, the availability of rodent tissues across the lifespan has provided a rich picture of EN diversity. At least 13 murine subtypes have been readily distinguished by immunohistochemistry, with 11 being observed in the myenteric plexus ^6^. Importantly, some mouse EN types exhibit similarities with those of humans ^1^.

EN with the most consistent morphology between humans and rodents are classified as Dogiel type I and II. Type I EN are motor neurons consisting of two distinct classes, including nitrergic inhibitory neurons and excitatory cholinergic neurons. Each subtype comprises ~5-10% of all ENs within the myenteric plexus. Type II neurons, also known as Intrinsic Primary Afferent Neurons (IPANs), are an important class of sensory neurons and interneurons that coordinate gastrointestinal motility and secretion, accounting for ~9% of all human myenteric neurons. IPANs have smooth cell bodies, several long, uniformly branching axons, and no dendrites ^7^. However, few selective molecular markers have been established for type I and type II ENs in mice ^6^ and even fewer are known for humans.

Other classes of human EN (type III – IX) and their corresponding marker genes are less consistent across species. For example, human type III neurons have one axon with many branched dendrites and are labeled by Calbindin 1 (*CALB1*). In rodents *Calb1* labels a subset of type II neurons [Brehmer, 2018] and the closest counterpart to human type III neurons are classified as “filamentous” ^2, 6^. Another example of cross-species differences, somatostatin (*Sst*) selectively labels filamentous neurons in mice, while in humans, *SST* labels type II neurons and other subtypes ^6^.

To bridge the gap in comparing analogous neuron subtypes between humans and other species, major efforts are needed to profile EN at single-cell resolution. Such endeavors have been hampered by the difficulty of isolating intact neurons from the intestine of adult mammalian species. EN are either sandwiched between the outer muscle layers of the gut wall or buried in the submucosa. Recent advances have been made in profiling pools of EN and glia using bulk sequencing from fetal mice ^8^, when neuronal processes are less-developed and cells survive tissue dissociation. Most recently, single-cell gene expression profiles of ~1100 ENs have been generated from postnatal mice around weaning ^9, 10^. Because neuronal differentiation is still ongoing at these stages, such profiles are unlikely to fully capture adult EN gene diversity^11^. Here, we profiled adult mouse EN and human myenteric ganglia from the duodenum, ileum, and colon. With these efforts, we sought to: 1-molecularly define myenteric neuron subclasses across multiple regions of the human and mouse intestine, 2-compare genes that mark neuron subclasses between humans and mice, and 3-identify segment-specific markers for neuronal subtypes. We discovered several previously uncharacterized EN subtypes in humans and identified regionally expressed EN markers conserved between humans and mice. The resulting atlas offers a foundation for future mechanistic studies of gene function and drug targeting, with tremendous potential to identify the etiology of ENS-related gastrointestinal diseases.

## Methods

### Animals

All experimental protocols were approved by the Institutional Animal Care and Use Committee at Vanderbilt University. Tg(Phox2b-HIST2H2BE/Cerulean)1Sout mice (MGI: 5013571), hereafter *Phox2b*-CFP, were bred to C57BL6/J and adult progeny of both sexes at 6-7.5 weeks of age were used. All mice were housed in a modified barrier facility on a 14-hour on, 10-hour off light cycle in high density caging (Lab Products Inc., #10025) with standard diet (Purina Diet #5L0D) and water *ad libitum.*

### Human tissue

This study was approved by the Vanderbilt University Institutional Review Board and classified as non-human subjects research. All tissue samples were received from post-mortem, de-identified healthy organ donors aged 18-35 years (Figure 1A). GI tissue harvest included duodenum (~8cm) beginning just distal to the pancreatic duct, ileum (~20cm) proximal to the appendix, and colon (~20cm) straddling the center of the transverse colon.

**Figure 1.**
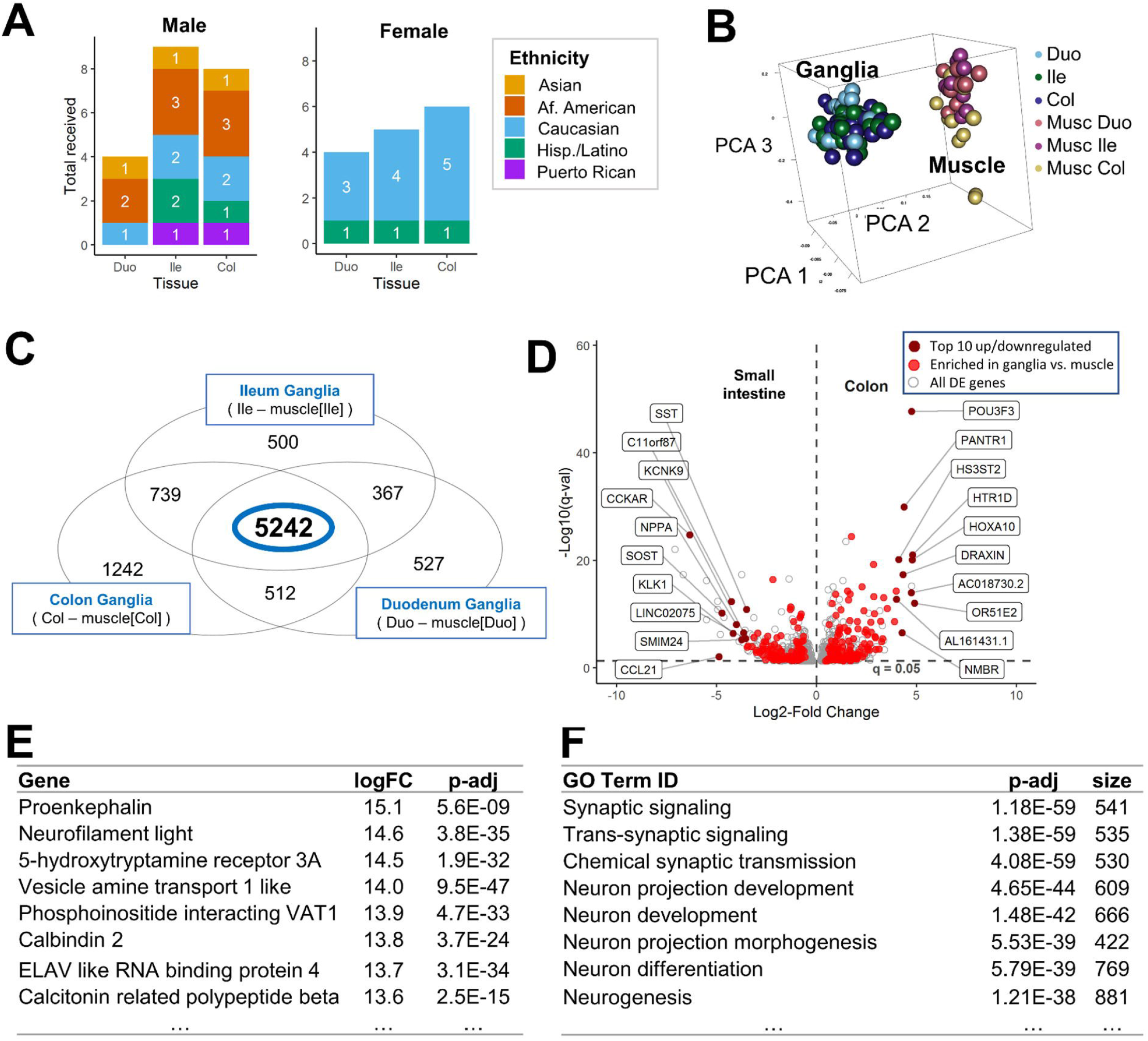
LCM RNA-Seq of healthy human enteric ganglia identifies commonly expressed genes across all intestine segments. (A) Demographic plot of intestinal tissue donors. (B) Human ganglia samples cluster distinctly from those of muscle in 3-D PCA space. (C) Venn Diagram illustrates differentially expressed genes between enteric ganglia and muscle when comparing expression between all intestine regions. (D) Volcano plot of differentially expressed genes between human colon and small intestine, with the top10 most up/downregulated genes annotated based on log_2_FC. (E) Most highly upregulated genes and GO terms (F) in myenteric ganglia relative to muscle, ranked by log_2_FC.

### Laser-Capture Microdissection (LCM)

Human tissue was sectioned at 10-μm and processed via LCM on an ArcturusXT™ for RNA as described ^12^. Samples with RNA integrity values greater than 6.8 were submitted for library construction and sequencing. In total, 111 ganglia samples, each (pooled from 1-3 LCM caps worth of ganglia) and 27 intestinal muscle samples were successfully sequenced.

### Preparation of single-nuclei suspensions

Mouse intestinal muscle laminar strips containing the myenteric plexus were peeled away from the submucosa on ice while submerged in DPBS with Mg^2+^ and Ca^2+^. Tissue was minced in ice-cold DPBS and pelleted by centrifugation. Nuclei were isolated using the NucleiEZ nuclei isolation kit (Sigma), with modifications described in Supplemental Methods. All steps were performed in a 4° cold room.

### Nuclei isolation from ENs

Nuclei were isolated by fluorescence-activated cell sorting (FACS) on a BD FACSAria III using a 100µm nozzle at 17 psi. Nuclei were first separated from cellular debris based on forward and side-scatter with doublet discrimination achieved by forward and side scatter pulse geometry gating. Neuronal nuclei were gated for 7AAD+ and high intensity of CFP from the *Phox2b*-CFP reporter (Fig. 2A, bright population)^13^.

**Figure 2.**
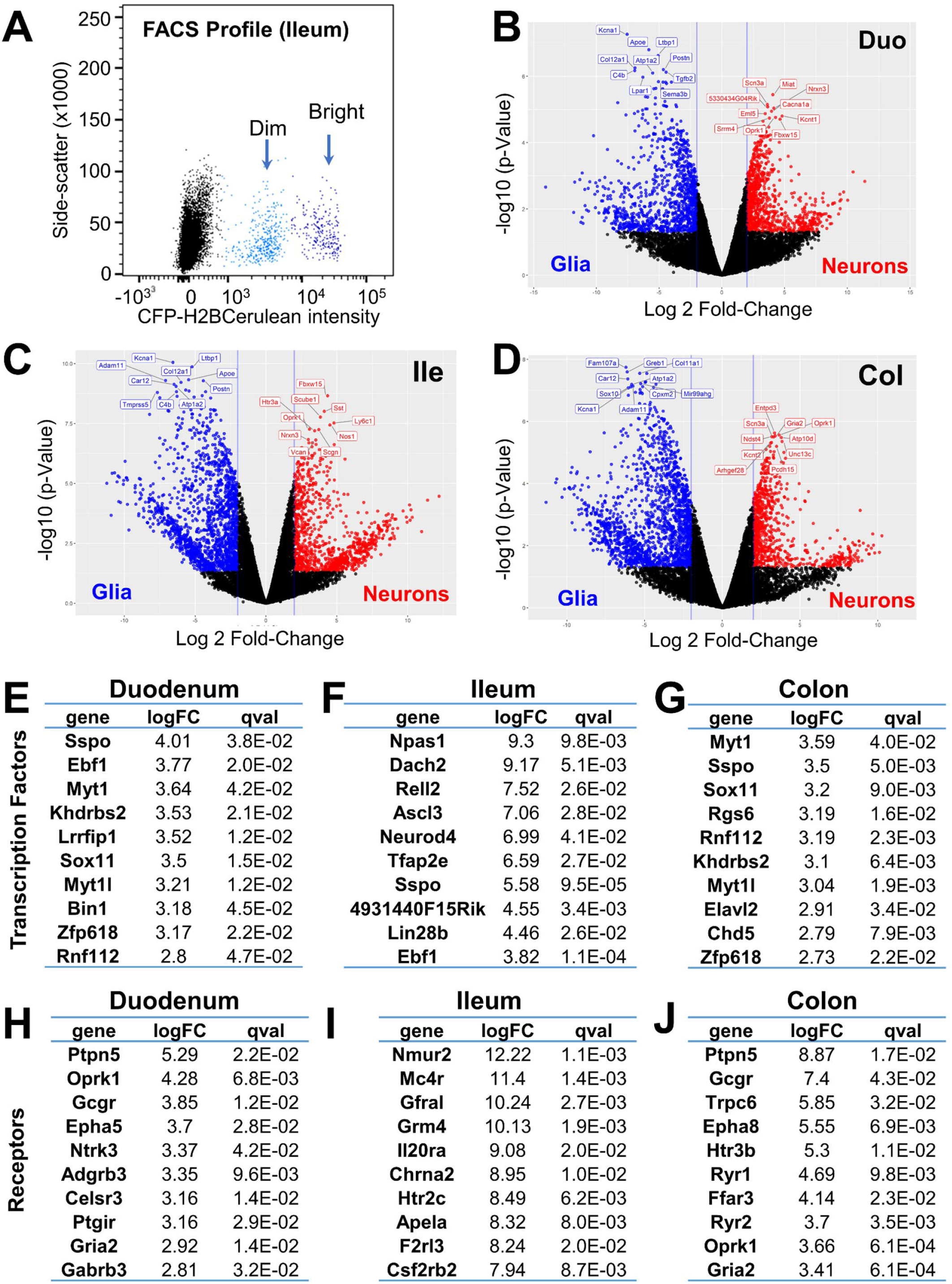
Neuron-specific genes identified via RNA-Seq of bulk adult mouse EN and glia. (A) FACS separation of neuronal (bright) and glial (dim) *Phox2b*-H2BCFP+ nuclei sorted for bulk RNA-Seq. (B-D) Volcano plots show differential gene expression in EN relative to enteric glia for each intestinal segment. (E-G) Top 10 most-upregulated transcription factors. (H-J) Top 10 most-upregulated receptors.

### snRNA-Seq encapsulation and sequencing

Single nuclei were encapsulated using both 10X Genomics and InDrop methods. For generation of 10X libraries, nuclei were encapsulated using version 3 Chromium Single Cell 3’ library reagents. Libraries were sequenced using a Nova-seq 6000 or Nextseq 500 and a paired-end 50 bp sequencing flow cell at a total depth of 3.6 Billion reads across nine 10X runs (Supplemental Figure 2A, n=9).

On the inDrop platform (1CellBio), nuclei were encapsulated and libraries were prepared using a modified version of the Cel-Seq method (See Supplemental Methods). Following sequencing, the combined total read depth from 8 samples amounted to 550 Million reads across all inDrop samples (Supplemental Figure 2A, n=6).

### Bulk RNA-Seq: cDNA library preparation and sequencing

Libraries were constructed from total human RNA or flow-sorted mouse nuclei using the Takara SMARTer kit per manufacturer’s protocol. Sequencing was performed on a HiSeq3000 or NovaSeq as 1×50bp or 1×100bp single-end reads, respectively. Human samples were sequenced to a mean depth of ~75 million reads with a mean total alignment rate of 98.12% for a total of over 11 billion reads. Eleven mouse EN and glia samples from the duodenum, ileum, and colon were sequenced, each having a mean depth of approximately 138 million reads and a mean alignment rate of 99.01%.

### Bulk RNA-Seq data processing

Base calls and demultiplexing were performed with Illumina’s bcl2fastq software. RNA-Seq reads were aligned to the Ensembl release 76 human or mouse assemblies with STAR version 2.0.4b. Gene counts were derived from all uniquely aligned, unambiguous reads by Subread:featureCount version 1.4.5. Isoform expression of known Ensembl transcripts were estimated with Sailfish version 0.6.13. Differential expression was performed using edgeR in conjunction with Limma-Voom. Full details in Supplemental Methods.

### snRNA-Seq data analysis

The sequencing output FASTQs were processed with CellRanger 3.0.2 using a modified mm10 reference enabling intron quantification to obtain a gene-cell data matrix. For inDrop, reads were filtered, sorted by their designated barcode, and aligned to the reference transcriptome (intron + exon) using DropEST pipeline (STAR). Mapped reads were quantified into UMI-filtered counts per gene. Raw matrix files were processed and analyzed using the R-package, Seurat (version 3) [Butler 2018]. The total number of snRNA-Seq runs merged from each intestinal segment included: Duodenum: 4, Ileum: 6, and Colon: 6. Data were batch-corrected and processed as described in Supplemental Methods. In total, 25,208 neuronal nuclei were successfully sequenced and passed quality control steps from a total of 16 snRNA-Seq runs.

### Fluorescence *in situ* hybridization and microscopy

*In situ* HCR version 3 was applied to visualize candidate markers for EN subtypes in human and mouse intestinal tissue using manufacturer protocols (https://www.molecularinstruments.com/). Probes were purchased from Molecular Instruments or were synthesized using the OligoMiner program [Beliveau, et al., 2018]. HCR was performed as described ^14^, using tissues samples from the duodenum, ileum, and colon of minimally three human donors. Before coverslipping, samples were treated with TrueBlack® dye to quench lipofuscin autofluorescence ^15^(See Supplemental Methods). Images were generated using a Leica DMI6000B or LSM880 confocal microscope, as indicated.

## Results

### Transcriptome catalog of human ENS genes from healthy young adult myenteric plexus

To capture baseline transcriptional profiles for total EN from healthy human intestine, we applied RNA-Seq to LCM material from myenteric ganglia and adjacent smooth muscle for both sexes and multiple ethnicities (Figure 1A). Tight clustering of sample replicates for EN and intestinal muscle with principal component analysis (PCA) indicated high data quality with absence of outliers (Figure 1B) and consistent patterns of gene expression within myenteric ganglia throughout the intestine. We compared transcriptional signatures from myenteric ganglia with those of adjacent smooth muscle in each bowel region to derive an EN-specific transcriptome catalog based on identifying genes expressed at significantly greater levels in myenteric ganglia relative to intestinal muscle (p-adj<0.05, Figure 1C). This approach detected 5,242 genes differentially expressed relative to muscle and stably expressed across the entire intestine (Figure 1D). Among this list, genes with the most highly up-regulated expression in the ENS included the well-known EN markers: *PENK, NF*, and *ELAVL4* (Figure 1E). Additionally, the most significant gene ontology (GO) biological functions for this gene list relate to the nervous system (Figure 1F). These results provide strong evidence we have successfully generated a comprehensive transcriptome catalog of the ENS from healthy human adults.

Segment-specific expression of genes enriched in myenteric ganglia was also detected for thousands of genes (Figure 1C). Separate analyses identified differential gene expression between the small intestine and colon for 254 genes, which also had selective expression in ganglia relative to muscle (Figure 1D).When examining differences in gene expression between the small intestine and colon, several genes showed substantial regional expression differences in ganglia, including *CCKAR*, *POU3F3,* and *HOX3A* suggesting potential region-specific ENS functions.

### Bulk RNA-Seq of Phox2b+ neuronal and glial nuclei identifies transcription factors and receptors expressed in ENs

To derive a comparable resource from adult mouse ENS for comparison with human ENS gene signatures, we collected nuclei from the intestines of *Phox2b*-CFP transgenic mice and sorted based on CFP intensity to separately isolate EN and glia (Figure 2A). Nuclear integrity and cell-type-specific gene expression were assessed by RT-PCR in FACS-purified nuclei pools, prior to RNA-Seq. Sorted nuclei were intact and exhibited excellent retention of gene signatures specific to EN or glia (Supplemental Figure 1). Pools of nuclei from ENs or glia were then sequenced from each segment of the intestine, leading to the identification of differentially expressed genes (Figure 2B-J). Some of the most highly up-regulated genes expressed in neurons from all segments included *Gcgr*, *Slc35d3*, *Scl26a4*, *Dnah11*, *Oprk1*, and *Sst*. Similarly, the most highly up-regulated glial genes across gut segments included *S100b*, *Prkca, Gpt2, Fads1, Ndrg1,* and *Limd1*. We estimate that ~40% of all genes enriched in neurons relative to glia are shared across all gut segments. This analysis also identified numerous transcription factors and receptors for each intestinal segment that are highly up-regulated in mouse EN (Figure 2E-J).

### snRNA-Seq from adult mouse duodenum, ileum, and colon identifies EN subtypes with some differences in regional prevalence

To assess the similarity of EN types in each intestinal region and relate cluster identity to markers expressed by known functional EN classes, we merged 10X snRNA-Seq datasets (see Supplemental Methods). Merged nuclei separated into 15 distinct clusters (Figure 3A) that exhibited expression of neuronal genes *Elavl4* and *Prpn* (Supplemental Figures 3 and 4B). A single cluster that lacked neuronal markers and exhibited glial character, consistent with known *Phox2b* expression in enteric glia ^13^, was removed during processing. Based on the gene lists derived from these remaining 15 remaining clusters, we assigned identities based on known immunohistochemical labeling of 11 functional myenteric neuron types ^6^ to each cluster. This approach allowed us to generate putative functional identities for 10 of the 15 snRNA-Seq clusters, including excitatory and inhibitory longitudinal muscle motor neurons, descending interneurons, and ascending interneurons (Supplemental Figure 3C-D). Notably, several of these EN subtypes appeared to be restricted to particular intestinal regions, based on the proportions of neurons derived from each bowel segment in clusters 7, 9, and 10 (Supplemental Figure 3A-B). Neurons in cluster 7 (*Chat/±Calb2/±Calcb*) and 10 (*Nos1+/Chat+*) mostly originated from the small intestine while cluster 9 (*Sst*+) had the greatest contribution of neurons from the colon.

**Figure 3.**
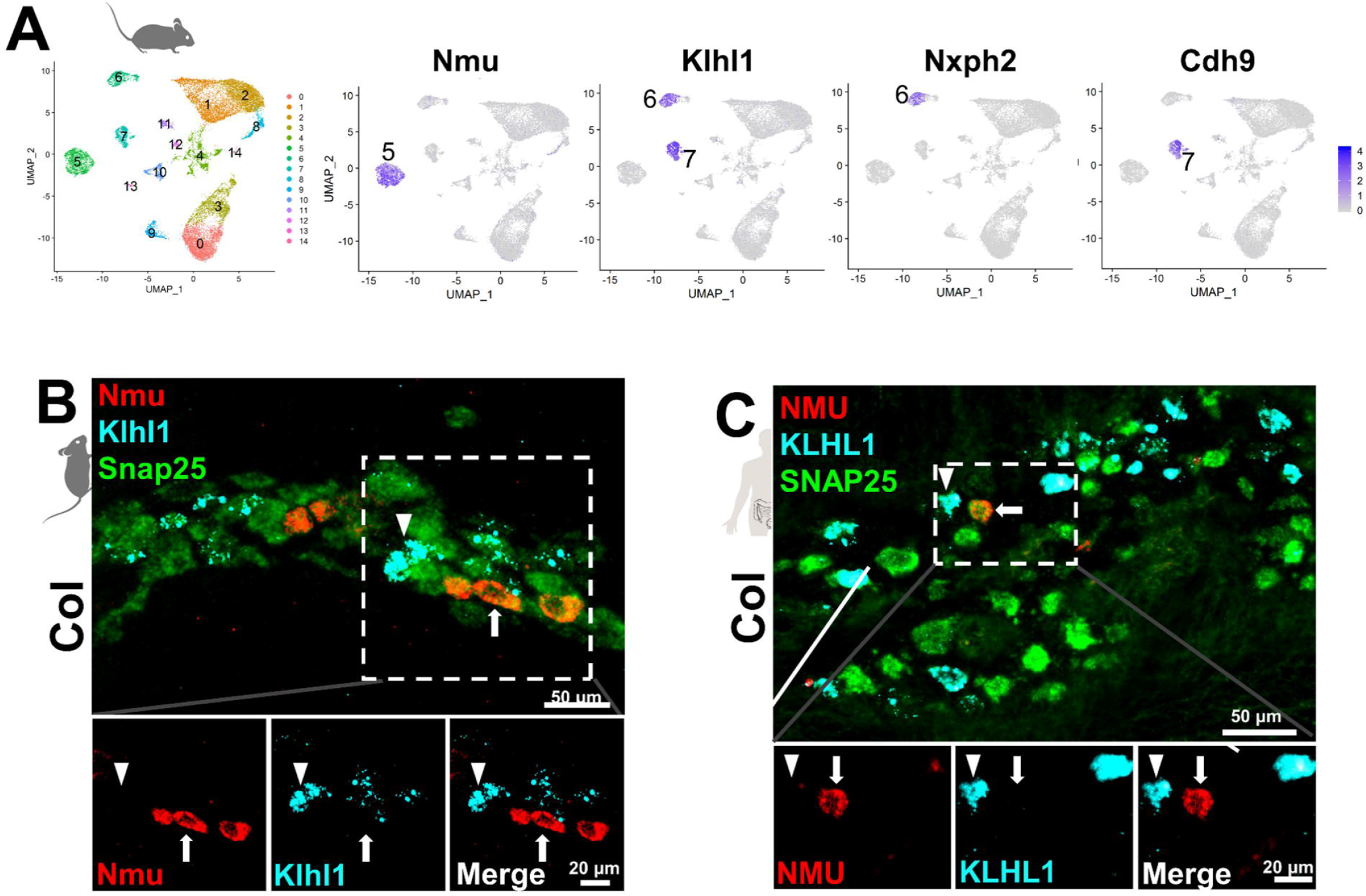
Genes marking neuron clusters in mouse identify human neuronal subtypes. (A) UMAP plot displays 15 distinct clusters detected in merged mouse snRNA-Seq data from all gastrointestinal regions with example genes marking discrete clusters. samples, showing the expression of each marker gene, arranged by segment. (B) *Nmu* and *Klhl1* expression visualized by HCR FISH mark distinct EN subtypes in adult mice and (C) Human myenteric plexus. Arrowheads mark Klhl1+neurons. Arrows indicate exclusively Nmu+ neurons. Snap25 (green, pan-neuronal).

We next examined classification of clusters that were not readily assigned to previously characterized EN subtypes, including clusters 4, 8, and 11-14. We considered the possibility that these clusters might be non-neuronal. However, KEGG functions examined for cluster-specific marker gene lists each contained multiple significant terms relating to the nervous system (Supplemental Figure 4A). The single-cell mouse cell atlas program (scMCA)^16^ also predicted that all clusters were neuronal with moderately high confidence, except for clusters 11 and 14, which exhibited strong predictions for mesenchymal and vascular/immune cell types (Supplemental Figure 4F) that may coincide with a recently reported mesenchymal lineage derived from the vagal neural crest ^17^. Further examination of these clusters showed robust expression of the early-neuronal genes *Elavl4* and *Prph* (Supplemental Figure 4B). Cluster 12 expressed *Sox10*, a gene that marks ENS progenitors; although the progenitor marker *Nestin*, was infrequently expressed in cluster 12 and appeared only sporadically in nuclei throughout all clusters (Supplemental Figure 4C-D). Markers for cycling cells (*Top2a,* Supplemental Figure 4E) were also sparsely observed throughout all clusters, with only moderate expression in cluster 13. We designate these clusters as functionally “unassigned” until further characterization can be pursued.

During our analysis of EN cluster gene expression, it became apparent that several clusters could be further subdivided, some of which were differentially distributed between intestinal regions. Specifically, clusters 5-7, 9, and 10 exhibited non-uniform expression of known EN subtype markers including *Calb1*, *Calb2*, and *Vip* (Supplemental Figure 5). This evidence suggests that these sub-clusters may represent up to eight distinct EN subtypes based on known EN markers^6^ (Supplemental Figure 5). Altogether, we estimate that there are up to 20 EN subtypes in the mouse myenteric plexus collectively across all intestinal regions.

### Selection of cluster-specific mouse EN subtype markers

To assess conservation of EN subtypes between species, we sought to identify cluster-specific marker genes for adult mouse EN that were also expressed in human EN. First, we selected highly expressed genes from the 15 main clusters detected among our mouse snRNA-Seq data for all bowel regions. We prioritized genes that were highly expressed in a single cluster, present within a high percentage of nuclei within that cluster, and minimally expressed in other EN clusters. From this gene set, we retained those that were highly expressed in human myenteric ganglia LCM/RNA-Seq data and minimally present in human intestinal muscle. This approach identified high-scoring markers for most of the mouse EN clusters (Supplemental Figure 3C. Notably, we observed that three exceptional EN clusters (5, 6 and 7) had many markers that were highly scored by the above criteria. These clusters were assigned an identity of intrinsic primary afferent neurons (IPANs) based on their expression of known mouse IPAN markers, including *Nefl*, *Calb2*, and *Calcb* (Supplemental Figure 3C-D) ^6^.

### Conservation of IPAN markers between humans and mice

Because IPANs are not well-characterized in healthy adult humans, we investigated the expression of the putative IPAN markers identified from mouse snRNA-Seq using HCR for *in situ* localization ^14^. As a prerequisite for generating neuron-specific marker genes as probes, we further excluded any cluster-specific genes with high expression in enteric glia and other cells of the bowel wall (Supplemental Methods). Multiple high-scoring marker genes remained for clusters 5, 6 and 7, including *Nmu* (cluster 5), *Klhl1* (6 and 7), *Nxph2* (cluster 6), and *Cdh9* (cluster 7). We applied HCR to visualize the most highly scored IPAN subtype markers, *Nmu* and *Klhl1*, in mouse and human intestinal tissue. Consistent with the cluster-specific expression seen in our mouse snRNA-Seq data, probes for *Nmu* and *Klhl1* labeled distinct EN subtypes in mice (Figure 3B), while orthologous *NMU* and *KLHL1* probes in human duodenum, ileum, and colon similarly labeled distinct EN (Figure 3C, Supplemental Figure 6C-D). We further applied HCR to detect additional genes co-expressed in cluster 5 (Nmu+) in mice, including *Dlx3* and *Otof* (Supplemental Figure 6). In both mice and humans, we documented coexpression of *DLX3* and *OTOF* in *NMU*+ EN (Supplemental Figure 6E-H). Our results illustrate similarities of gene expression for this subtype in mice and humans; however, we noted that *OTOF* and *DLX3* are more broadly expressed, being present in *NMU*-negative EN of humans (Supplemental Figure 6F,H).

### *Nmu* and *Klhl1* mark IPANs in mice, while in humans, only *NMU* labels IPANs

Among rodents, *Nmu* is known to label guinea pig IPANs and a recent study has confirmed this in mice based on co-localization with the established IPAN marker *Calb2* ^6, 10^. Our finding of *Nmu* in mouse IPANs is consistent with these studies, although to our knowledge *NMU* has been untested in the human ENS. Further, we also identified *Klhl1* as a novel marker of IPANs in mice. We established that *Klhl1* labels type II neurons (IPANs) in the myenteric plexus of mice using *Nefl* for co-labeling ^6^ (Figures 5A-B) and documented strong, localized expression of *Klhl1* within *Nefl*+ EN. An IPAN identity for *Klhl1*+ (cluster 6) myenteric neurons in mice is further supported by a recent report of type II morphology and connectivity for *Klhl1*+ neurons ^10^. We subsequently assessed the expression of *NMU* and *KLHL1* in human myenteric ganglia. Human IPANs are defined by co-expression of Somatostatin (*SST*) and *CALB2*, although each marker labels many other EN types ^18^. We show that *NMU* labels duodenal, ileal, and colonic human IPANs (Figure 4, Supplemental Figure 7). Approximately 50-70% of human IPANs co-expressed *NMU* in the small intestine, with slight variations between donors and depending on the position of the tissue section within the ganglia.

**Figure 4.**
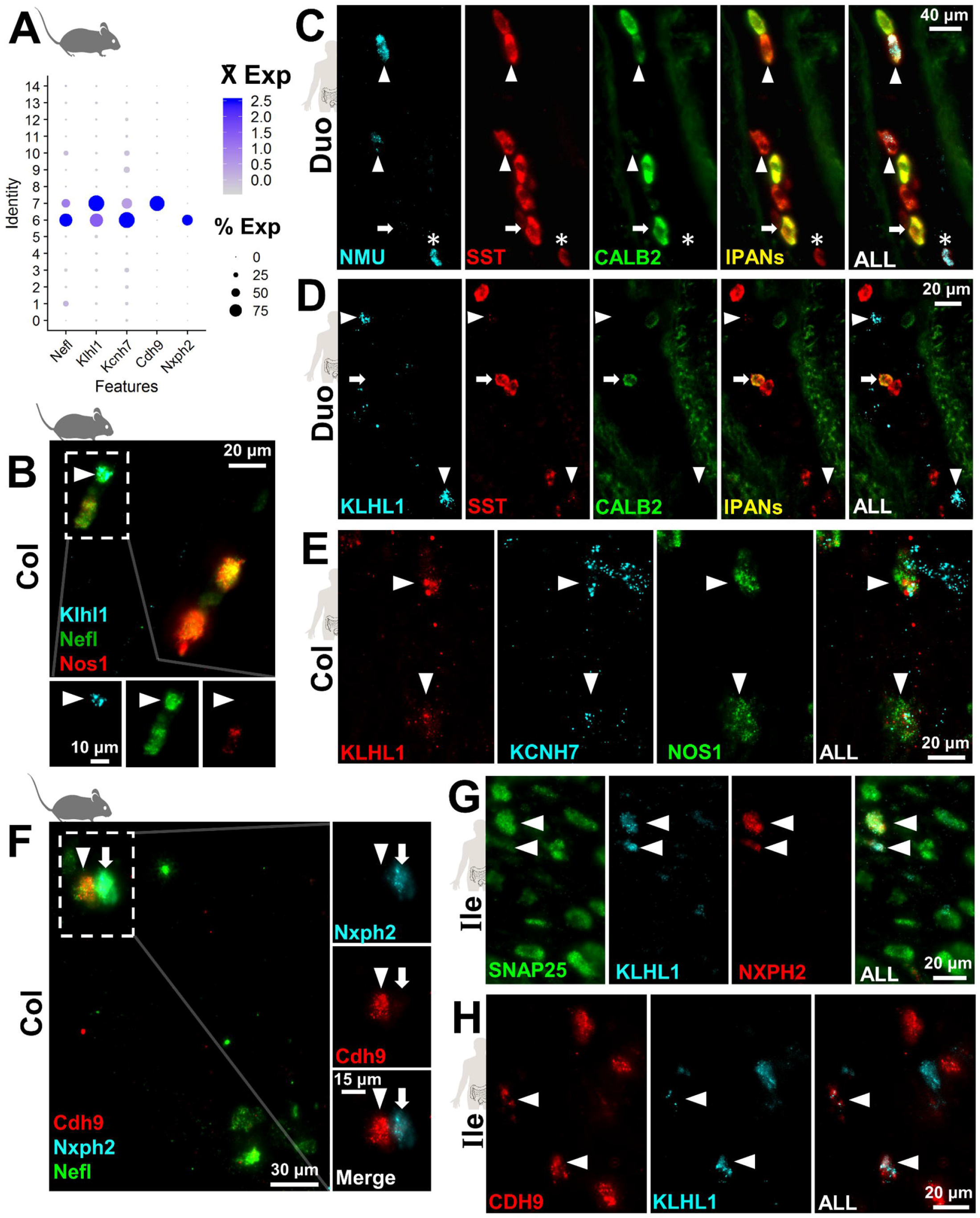
Mouse IPAN markers label multiple neuron types in human ENS. (A) Expression of mouse IPAN marker *Nefl* marker in snRNA-Seq dot plot labels clusters 6 and 7 concurrent with subtype marker genes *Klhl1, Kcnh7*, *Nxph2,* and *Cdh9*. (B) HCR-FISH confirms co-expression of *Klhl1* with *Nefl*+ IPANs (arrowhead) in mice. (C) Human *NMU* expression localizes with many IPANs (SST+/*CALB2*+; arrowheads) but not all (arrow). Some human *NMU*+ neurons are not IPANs (asterisk). (D) Human *KLHL1* expression (arrowheads) is distinct from IPANs (SST+/*CALB2*+, arrow). (E) *KCNH7* colocalizes with many *KLHL1*+ neurons (arrowheads), including some NOS1+ neurons. (F) HCR-FISH confirms *Cdh9* (arrowheads) and *Nxph2* (arrows) are expressed in distinct IPAN subtypes. (G) *NXPH2* is co-expressed with *KLHL1*+ ENs (arrowheads) in human ileum. (H) *CDH9* is expressed in *KLHL1*+ neurons (arrowheads) and some *KLHL1*[-] neurons in human ileum.

**Figure 5.**
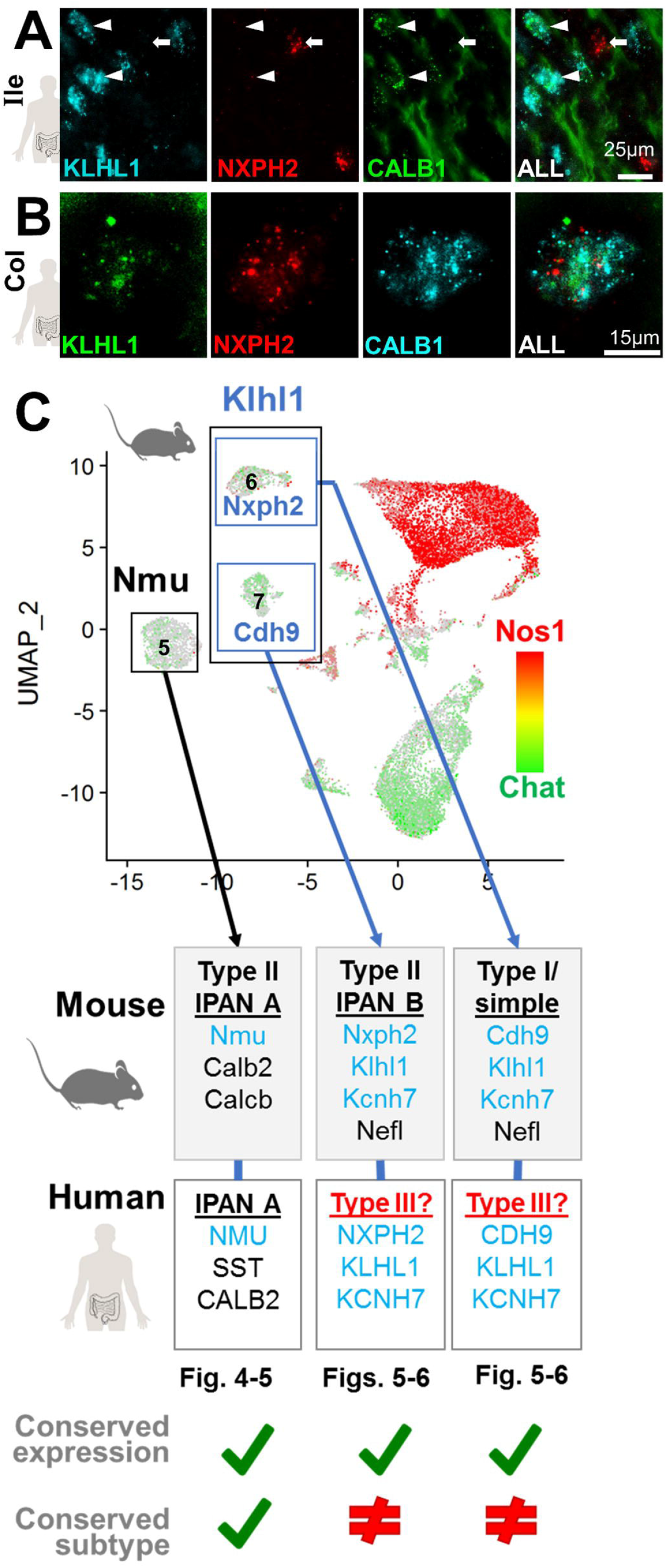
Dichotomous expression of mouse IPAN markers compared to human ENS. (A) In human ileum *CALB1*+ (type III neuron marker) co-localizes with *KLHL1* (arrowheads). Contrasting with mouse, *NXPH2* rarely localized with *KLHL1* or *CALB1* (arrow). (B) Co-expression of *CALB1*, *KLHL1*, and *NXPH2* marks a distinct human EN subtype in colon. (C) Summary of the cross-species comparisons for subtypes of ENs examined here.

Unexpectedly, fewer IPANs expressed *NMU* in the colon (~5-10%, Supplemental Figure 7), leading us to anticipate that the remaining *NMU*[-] IPANs would express *KLHL1*.

However, despite repeated attempts, we did not detect *KLHL1* expression in any IPANs along human intestine based on *SST*/*CALB2* labeling (Figure 4D, Supplemental Figure 7C-D).

### *KLHL1* labels a discrete, non-IPAN subtype among human myenteric neurons

We further investigated whether *KLHL1* is part of a conserved set of subtype marker genes in human IPANs, or whether it labels completely different EN subtypes across species, like *CALB1* ^1, 6^. To this end, we examined the expression of KCNH7 in *KLHL1*+ neurons of the human intestine, because *Kcnh7* is co-expressed with *Klhl1* in EN clusters 6 and 7 of mice. *In situ* HCR labeling showed that *KLHL1* and KCNH7 are co-expressed in a discrete population of myenteric neurons (Figure 4E). Although *Klhl1* and *Kcnh7* were highly co-incident in mice based on snRNA-Seq data, expression of *KLHL1* and KCNH7 in human myenteric neurons did not completely co-localize within EN and were observed in a much larger proportion of EN than in mice. We further examined whether KCNH7 might label human IPANs more effectively than *KLHL1*, but similarly did not identify any KCNH7 expression in IPANs based on labeling with *SST*/*CALB2* (Supplemental Figure 7E). These observations indicate that although *KLHL1* and KCNH7 do not label human IPANs, they mark a completely distinct EN subtype.

Given the coincident expression of *KLHL1* and KCNH7, we further evaluated the novel *KLHL1*+ subtype(s) among human EN. In mice, *Nxph2* and *Cdh9* discretely labeled *Klhl1*+ neurons of clusters 6 and 7, respectively (Supplemental Figure 7F-G), and co-localized with the mouse IPAN marker, *Nefl*, by HCR(Figure 4F). Consistent with our findings in mice, *CDH9* and *NXPH2* labeled human *KLHL1*+ neurons throughout all intestinal regions (Figures 5G-H, Supplemental Figures 7H-J, 8A-C). However, *CDH9* and *NXPH2* were also expressed in human EN lacking *KLHL1* expression and some specimens exhibited colocalization of *CDH9* and *NXPH2* in contrast to labeling distinct subtypes in mice (Supplemental Figure 7). *KLHL1*+ neurons in humans also appeared to be expressed in a variety of neuron types with a wide range of NOS1 expression, in contrast to *Klhl1*+ ENs in mice that are mostly *Nos*-negative (Supplemental Figure 7). Overall, we conclude that *CDH9* and *NXPH2* are conserved markers for subtypes of *KLHL1*+ neurons between species. While we observed similar numbers of these neurons, *CDH9* and *NXPH2* do not exclusively mark human *KLHL1*+ neurons as in mice.

To clarify the cellular identity of human *KLHL1*+ ENs, we examined the possibility that *KLHL1* expression labels type III neurons. In humans, type III neurons are observed in the small intestine and are strongly labeled by *CALB1*. However, *CALB1* is not entirely exclusive to human type III neurons, as a small proportion of type II and sporadic type I ‘spiny’ and ‘stubby’ neurons also stain for *CALB1* ^2^. We observed the majority of *CALB1*+ neurons in the duodenum and ileum strongly express *KLHL1* (Figure 5, Supplemental Figure 8), suggesting that *KLHL1* is a type III neuron marker in the small intestine. Prior work has shown that *CALB1* labels a yet-unclassified subtype of EN in the human colon ^2^. Using HCR, we consistently found that the majority of *CALB1*+ neurons in human colon expressed *KLHL1*+ (Figure 5B) and that *NXPH2* was co-expressed in all *CALB1*+ neurons of human colon, analogous to mice (Figure 5).

However, unlike mice, *CDH9* expression was not observed in *KLHL1*+/*CALB1*+ neurons of the human colon (Supplemental Figures 7G, 8C). In contrast to prior reports, we detected fewer *CALB1*+ neurons in the colon with HCR, compared with prior *CALB1* antibody labeling in human colon ^2^, which might be attributable to the high stringency of HCR relative to the potential cross-reactivity with antibodies. We illustrate our findings of IPAN subtype marker conservation in mice and humans in Figure 5C. Type II ENs in mouse cluster 5 (*Nmu+*) and in humans appear to both share similar morphology ^1, 6^ and selective marker genes. However, mouse clusters 6 and 7 were not morphologically conserved ^1, 6, 10^, despite sharing similar markers genes across species (*KLHL1*, *KCNH7, NXPH2*, and *CDH9*).

### Regionally expressed neuronal subtype markers in humans and mice

Our comprehensive study design allowed us to examine which neuron subtypes are present along the entirety of the intestine and whether any subtypes are found only in particular intestinal regions. Moreover, we were able to assess whether EN subtypes present throughout the entirety of the intestine can exhibit graded expression of distinct marker genes in different bowel regions. This possibility was first confirmed using our mouse snRNA-Seq data, which revealed multiple subtype-specific genes have regional expression patterns (Supplemental Figure 9). The presence of regionally-expressed genes in mouse EN subtypes raised the possibility that human EN subtypes could share similar region-specific expression. We subsequently identified regionally expressed genes in human myenteric ganglia that were expressed in mouse EN subtypes (Supplemental Methods, Figure 7A-C), including *CCKAR*, *RBP4*, *WIF1*, *SYT15*, *POU3F3*, and *NEUROD6*. However, most regional expression patterns of genes were not shared between species.

*CCKAR* was examined further as a result of its marked expression differences in the small and large intestine of both mice and humans (Figure 6D-F). In mice, *Cckar* expression was restricted to clusters 6 and 3 (Figures 7D-E, Supplemental Figure 10A, Supplemental Figure 11). We examined *Cckar* expression in mouse cluster 6 subtype, given its classification as a putative IPAN, based on *Nefl* expression. *Cckar* expression was coincident with *Nefl* in mouse intestinal tissue using HCR and nearly entirely colocalized with *Nxph2* and *Klhl1* (Figure 6G, Supplemental Figure 10B). Far fewer *Cckar*+*/Klhl1*+ neurons were found in the ileum and colon (Figure 6H-I), consistent with snRNA-Seq data indicating that only 1% of neurons in the ileum and colon cluster 6 express *Cckar*.

**Figure 6.**
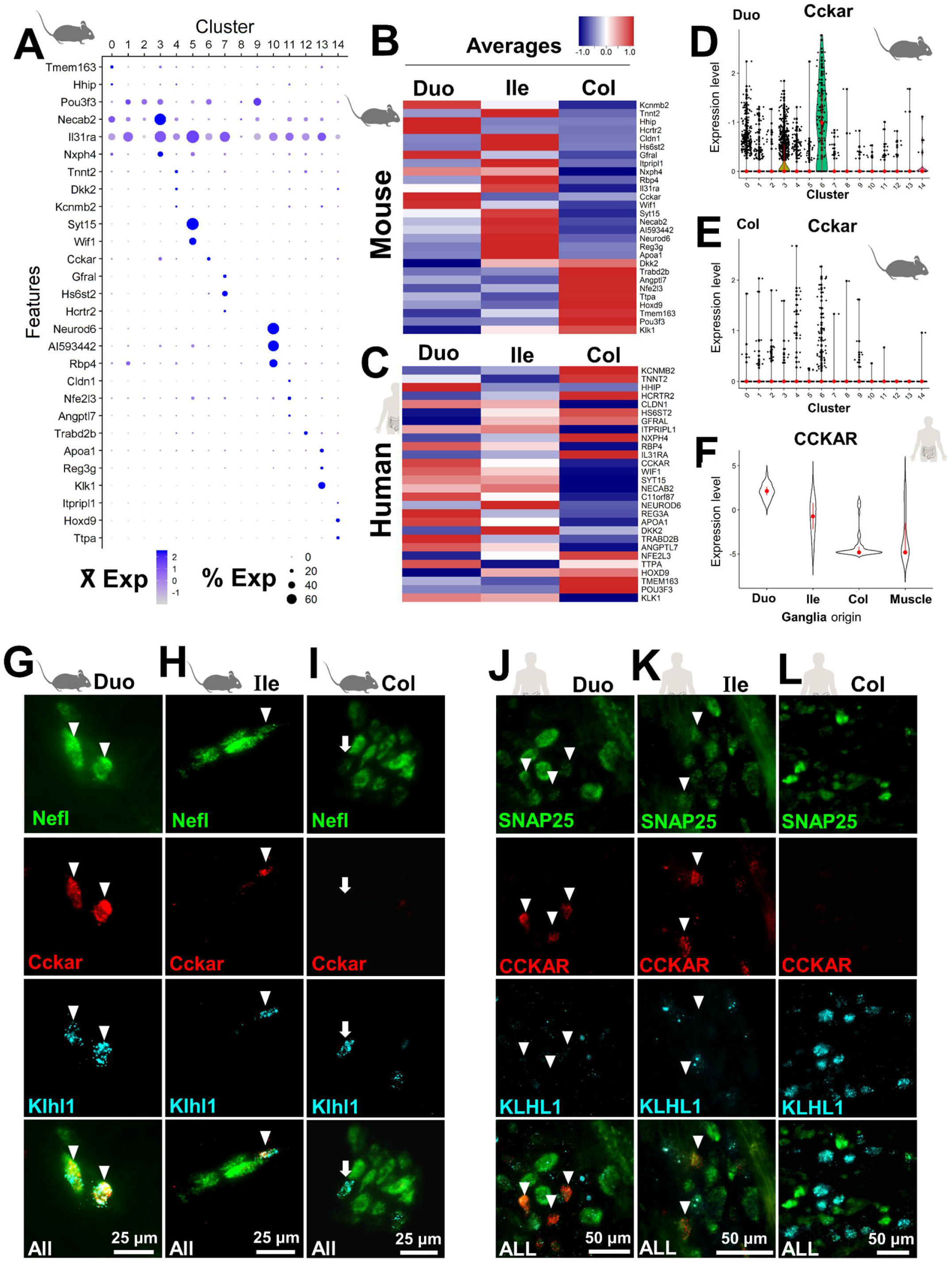
Cross-species comparison of regionally expressed genes in myenteric neuron subtypes. (A) Dot plot illustrating regionally expressed genes for each cluster from mouse snRNA-Seq data merged from all intestinal segments. (B) Heatmap displaying mouse genes from A ordered by the segment with highest expression. (C) Heatmap of human ortholog expression from myenteric ganglia. (D-E) Violin Plots depict the differences of *Cckar* regional expression for cluster 6 neurons. (F) Violin Plot of human *CCKAR* expression shows similar regional expression. (G-I) HCR FISH images for mouse *Cckar* reveals frequent *Nefl* and *Klhl1* co-localization in duodenal IPANs (arrowheads; cluster 6 in A); infrequent co-expression in the ileum and colon, and absence of *Cckar* in some colonic IPANs (arrows). (J-L) Human HCR FISH images show that mice and humans have similar regional expression of *CCKAR* but that expression is not localized to *KLHL1*+ neurons in humans (arrowheads), in contrast with mice.

Given that frequent, high *Cckar* expression among duodenal neurons was generally absent from other segments, we further examined snRNA-Seq data to assess whether *Cckar* expression defines distinct EN subtypes in cluster 6. Notably, the distribution of *Cckar*+ neurons for the duodenum, ileum and colon in UMAP space was uniform and intermingled with most *Cckar*[-] neurons, (Supplemental Figure 11), suggesting that most *Cckar*[-] neurons in the ileum and colon are not distinct from other *Cckar*+ neurons in this cluster. We conclude that most neurons in cluster 6 likely represent a single EN subtype distributed throughout the intestine with regional differences in gene expression, as illustrated in Supplemental Figure 11C.

*CCKAR* expression in humans was robust within myenteric neurons of the duodenum, as assessed with HCR, although few *CCKAR*+ neurons were observed in the ileum and none were detected in the colon (Figures 7J-L). Unexpectedly, *CCKAR* expression was not localized to any neurons expressing *KLHL1* (Figure 6J-L), in contrast to mice (Figure 6G-I). Attempting to determine the identity of *CCKAR*+ neurons, we examined the co-expression of *CCKAR* with alternative neuronal subtype markers. *CCKAR* expression was not detected in any IPANs, despite observing many instances of co-expression with many *OTOF* (Supplemental Figure 10C). Based on these findings, it does not appear that human *CCKAR* is expressed in many more neuronal subtypes than in mice, despite both species exhibiting robust and region-specific expression within the ENS.

## DISCUSSION

Here, we present a transcriptional catalog of genes expressed in healthy adult human ENS that is complemented by the parallel generation of an atlas from EN and enteric glia of adult mice derived from pooled “bulk” populations and snRNA-Seq. The resulting atlases provide a means of localizing specific neuron types *in situ* and have tremendous potential to identify deficits in the ENS that contribute to GI disease.

We identified novel neuronal subtypes that are conserved between humans and mice. Prominently, *NMU*+ expression identifies IPANs in both species based on coincident labeling with known IPAN markers (*Calb2/Calcb* in mice; *CALB2/SST* in humans). Similarly, we document conserved expression of *NXPH2* in human and mouse colon although the functional neuron subtype marked by this gene remains to be determined. Other genes marking EN subtypes were divergent between humans and mice, based on the subtype-specific marker genes we investigated. Specifically, *Klhl1* marked IPANs in mice, but not in humans. Instead, *KLHL1* labeled human Type III neurons based on colocalization with *CALB1*. Despite many similarities in EN subtype marker gene expression between the mouse and human, our findings suggest that caution is needed when making cross-species inferences for specific EN subtypes.

Our strategy to examine multiple regions of the GI tract with snRNA-Seq identified a total of up to 20 myenteric EN subtypes commonly present throughout the entire intestine. We also obtained strong evidence that many EN subtypes have distinct, regional expression patterns of select marker genes in the intestine of both humans and mice. In humans, *NMU* expression labels the majority of small intestine IPANs (50-75% of total IPANs) with higher prevalence than any marker previously reported. However, *NMU* only labeled 5-10% of human colonic IPANs (Supplemental Figure 7B), consistent with the lower level of expression of colonic *NMU* in our human LCM RNA-Seq data (data not shown). The differential expression of *NMU* in EN along the GI tract illustrates the regional expression of a subtype marker gene in humans and also implies heterogeneity of human colonic IPANs. Similarly, we demonstrated with snRNA-Seq and HCR that *Cckar* was present at high levels in 90% of duodenal *Klhl1*+ neurons in mice; although, it was detected in only 1% of *Klhl1*+ ileal and colonic neurons (Supplemental Figure 11). Similar regional expression of *CCKAR* was observed in human EN, with many *CCKAR*+ neurons observed in the duodenum, fewer in the ileum, and none in the colon (Figure 6). Multiple instances of regional marker gene expression are evident in our mouse snRNA-Seq data, with *Otof* (cluster 5), *Grp* and *Calcrl* (cluster 7), and *Grp* (cluster 9) exhibiting prominent expression differences among duodenum, ileum, and colon (Supplemental Figure 5). We also show several differences in prevalence of mouse EN subtypes, with some clusters being derived mostly from the small intestine or colon (Supplemental Figure 3). Collectively, our observations of regionally expressed subtype-specific genes raises the possibility of developing therapeutics to target specific EN subtypes in one intestinal region while leaving neurons in other regions of the gut unaffected.

Analysis of IPANs is of critical interest given the prominent role played by this subtype in the ENS. Our findings confirm the existence of multiple IPAN subtypes previously indicated for mice ^6, 10^ and expand the marker toolkit available to localize these EN. The potential existence of multiple types of IPANs in humans is open to further investigation. A recent pre-print on BioRxiv applied a ground-breaking method to capture colonic EN from colon cancer patients for snRNA-Seq ^19^. Based on this initial small dataset, the authors concluded there is likely only one subtype of human colonic IPAN. Our observation that *NMU* is expressed in only a small subset of human colonic IPANs suggests there may be more than one IPAN subtype. Additional snRNA-Seq isolations from human tissue focusing on this population could prove helpful in generating a more complete picture of IPANs in the adult ENS.

Our study significantly expands knowledge regarding the regional expression of EN subtype-marker genes in the human gastrointestinal tract and illustrates the ability to detect EN subtypes independent of immunoreagents. Spatial distribution patterns of neurons throughout the gastrointestinal tract of adult humans were previously limited to only a few subtypes, including type III EN restricted to the small intestine and “giant” type II neurons of the upper duodenum, and were also limited to diseased intestinal tissue ^1, 2^. By targeting collection of healthy gastrointestinal tract tissues from young individuals (18-35 years), our study establishes a baseline for gene expression and spatial distribution of EN subtypes. These data will inform future efforts to identify the underlying etiology of gastrointestinal diseases, such as chronic constipation, geriatric fecal impaction, and inflammatory bowel disease, which are associated with a loss of EN over time. Mapping EN subtypes in aged individuals is a future need that our data will facilitate.

Efforts are ongoing to derive EN from induced pluripotent human stem cells ^20^. While stem cell transplantation studies have successfully generated ENs that integrate into aganglionic bowel ^21, 22^, it has not been possible to assess whether integrated cells accurately reflected adult EN subtypes. Our profiling of adult EN provides the first benchmark resource of its kind that can be used to assess whether EN generated by directed differentiation mimic their *in vivo* counterparts.

In summary, our study is the first to undertake a comparative analysis of EN subtypes between species, across multiple intestinal regions. This effort identified EN classes present all along the intestine with identification of marker genes and EN subtypes that exhibit regionality. Our application of HCR has permitted unprecedented visualization of EN subtypes without antibody limitations and an initial mapping of human EN subtypes across multiple bowel segments. The resources of this study offer both an improved framework for diagnosis of enteric neuropathies and other GI diseases with a neuronal basis.

## Supporting information

Supplementary Figure Compilation

## Abbreviations

CALB1: Calbindin 1
CALB2: Calbindin 2
CCKAR: CCK Receptor Type A
CDH9: Cadherin 2
CHAT: Choline acetyltransferase
EN: Enteric neuron
ENS: Enteric nervous system
FACS: fluorescence-activated cell sorting
FISH: fluorescence *in situ* hybridization
GO: Gene Ontology
HCR: Hybridization chain reaction
IPAN: Intrinsic Primary Afferent Neuron
LCM: Laser-Capture Microdissection
NEFL: Neurofilament
NMU: Neuromedin U
NOS1: Nitric oxide synthase
NXPH2: Neurexophilin 2
PCA: Principal Components Analysis
SNAP25: Synaptosome Associated Protein 25
snRNA-Seq: single-nucleus RNA-Sequencing
Sox10: SRY (sex determining region Y)-box transcription factor 10
SST: Somatostatin

## Acknowledgements

We are grateful to the organ donors, their families, and staff of the International Institute for the Advancement of Medicine and Tennessee Donor Services. Technical advice was provided by Drs. Moustafa Attar, Seoeun Lee, Lori Zeltzer, and Bob Matthews.

## Supplemental Figure Legends

**Supplemental Figure 1.** Isolated nuclei from ENs and glia retain RNA signatures of cellular identity

(A) Nuclei isolated from the “Bright” population of myenteric ENs from Phox2b-CFP(H2B) mice retain cerulean fluorescent protein (CFP) fluorescence throughout the isolation procedure and FACS.

(B) Gene expression of nuclei from EN (Bright) and glia (Dim) isolated by FACS from adult Phox2b-CFP mice. Expression was assessed using RT-PCR for nuclear RNA isolates from the duodenum (Duo), ileum (Ile), and colon (Col). Expression of an EN marker, Synaptotagmin 2 (*Syt2*), was detected only in the Bright population of FACS-purified nuclei and was not detected in nuclei from the Dim population. Conversely, Forkhead box D3 (*FoxD3*) was detected in nuclei from the Dim, but not Bright nuclei populations. Importin 8 (*Ipo8*) served as a loading control and indicated even loading of the PCR in each lane of the gel. “No-RT Control” = negative control with no reverse-transcriptase added to the RNA; “Whole Ile” = whole ileum sample, which contains at least a small amount of RNA from both neurons and glia; “gDNA” = mouse genomic DNA.

**Supplemental Figure 2.** SnRNA-Seq run information and batch integration.

Summary of all snRNA-Seq runs using the 10x and inDrop platforms, indicating the total runs for each gut segment, total reads, and the final numbers of nuclei before and after filtering. **\***Usable nuclei were identified after eliminating empty droplets and doublets as described in the Supplemental Methods.

**Supplemental Figure 3.** Identification of putative myenteric EN subtype identities in mice and their distribution along the full length of the intestine.

(A) Split-out UMAP plots of batch-corrected nuclei after merging runs from all gut segments using only 10x data (Col=Colon, Duod=Duodenum, Ile=Ileum).

A relatively homogenous distribution of neurons was observed for each cluster across all 10x runs, although there were proportionally fewer nuclei from the colon observed in clusters 7 and 10 and fewer nuclei observed in the duodenum for cluster 9.

(B) The composition of nuclei from each gut segment are listed for all clusters, with significant differences in the contribution of nuclei from each gut segment highlighted in blue.

(C) Putative cluster identities are proposed for each mouse snRNA-Seq cluster, based on recognized EN subtype markers from literature (“Known Markers”). Cluster identities that could not be confirmed in literature are listed as “Unassigned.” The 3 highest-scoring markers from each cluster are listed for mouse data (“Top 3 Mouse”), alone, as well as for the markers having human gene orthologs that were significantly enriched in human myenteric ganglia (“Top 3 Human-Match”).

(D) Proposed EN subtype identities for mouse snRNA-Seq clusters listed in panel C (“Proposed Subtype Identity”) are overlaid onto the original UMAP plot.

**Supplemental Figure 4.** Classification of putative subtype identities for mouse snRNA-Seq clusters not identified by published EN subtype markers.

(A) KEGG functions were derived from marker gene lists for each cluster. Here, the most significant nervous system-related terms are listed for clusters 4, 8, and 11-14, which had unassigned cluster identities based on published EN subtype markers. The majority of clusters, aside from 8 and 12, had a Bonferroni-adjusted p-value of < 0.05.

(B) To further investigate the identities of unclassified EN clusters, we examined the expression of early-neuronal commitment genes, *Elavl4* and *Prph*. *Elavl4* was expressed robustly in all clusters but had lower expression in a portion of nuclei from the unclassified clusters 4,8, and 11-14. *Prph* was similarly expressed at robust levels in the majority of nuclei, although expression levels varied quite a bit between clusters.

(C) Expression of glial marker gene, *Sox10* was observed only in cluster 12. This cluster was not excluded along with a separate glia cluster during pre-screening, because it strongly co-expressed pan-neuronal markers.

(D) The expression of the neural progenitor marker, Nestin (*Nes*) was assessed and appeared to be sporadically expressed in nuclei of all clusters. However, most cells in each cluster had no *Nes* expression, including cluster 12.

(E) Dividing cells, marked by expression of *Top2a,* were observed only sporadically in most clusters.

(F) Predicted cell types were determined with the single-cell mouse cell atlas (scMCA) program, by inputting the averaged expression levels of genes from all cells in each cluster. Neuronal identity was predicted for most clusters, other than 11 and 14, with slightly lower confidence being given to cluster 13. The top cell characteristic prediction for cluster 11 was mesenchymal/epithelial and cluster 14 was mostly endothelial. Notably, clusters 8 and 12, which did not have significant KEGG functions relating to the nervous system in panel A, had highest predictions for neuronal identity.

(G) Examination of several vascular and immune cell markers indicated robust expression of several endothelial and mesenchymal markers in cluster 14. Clusters 11-13 exhibited expression of immune cell markers in a small subset of nuclei.

**Supplemental Figure 5.** Subclustering of mouse snRNA-Seq EN subtypes identifies putative EN subtypes with differences in their regional distribution throughout the intestine.

(A) Cluster 5 can be divided into two subclusters in the small intestine and only one cluster in the colon, based on marker gene expression. *Calb2* and *Otof* are expressed in nuclei at the bottom of the cluster in the small intestine, although these markers did not subdivide the colon cluster by its marker gene expression, which was present in all neurons for *Calb2* but virtually not expressed in the case of *Otof*. Additionally, *Rab27b* was mostly restricted to the upper subcluster and was expressed in most nuclei in the colon for this cluster.

(B) Cluster 6 contained 2 distinct subclusters, the smaller of which was distinguished by the expression of many marker genes, including *Calb1* and *Rfx6*.

(C) Cluster 7 appeared to contain a total of at least 3 distinct subclusters, one of which was completely segregated from the main cluster, and was marked by discrete expression of *Carmn, Chrm2*, and several other markers. Additionally, discrete expression of *Grp* and *Calcrl* was noted in the left side of the main cluster. Few neurons were contributed to all subclusters by the colon, indicating this subtype is largely restricted to the small intestine.

(D) Cluster 9 segregated into two distinct subclusters in all segments. However, relatively few neurons derived from the duodenum were observed in this cluster, indicating this main putative subtype regionally-restricted to the distal bowel. The expression of *Grp, Calcb,* and *Hoxd5* were preferentially localized to the upper subcluster of the colon, but nuclei of the duodenum and ileum did not share the same pattern of gene expression, indicating they may only represent a single subtype.

(E) Cluster 10 appeared to contain 3 distinct subclusters based on discrete marker gene expression. The colon had very few neurons, proportionally, most of which were restricted to the bottommost cluster. *Vip* and *Ptprm* expression was mostly observed in neurons of the bottommost subcluster, with some expression apparent in neurons extending towards the uppermost portion of the main cluster. Expression of *Ednra* and *Npas3* were observed only in nuclei towards the top of the cluster. *Rigs4* and *Gm1673* were discretely expressed in the leftmost subcluster.

**Supplemental Figure 6.** Additional analysis of myenteric EN subtypes that are conserved between adult humans and mice.

(A-B) SnRNA-Seq expression plots from mouse ENs illustrate the selective expression of *Dlx3* and *Otof* cluster 5.

(C-D) HCR FISH images of *NMU* and *KLHL1* expression in distinct neuronal subtypes of the ileum (C) and duodenum (D). In both panels, arrowheads mark *NMU*+ neurons and arrows mark *KLHL1*+ neurons.

(E-H) HCR FISH images demonstrating consistencies between subtype markers across species for markers identified in cluster 5 from mouse snRNA-Seq.

(E) *Otof* and *Dlx3* are selectively co-expressed in mouse EN of the ileum in similar frequencies observed in snRNA-Seq data. Arrowheads indicate instances of colocalization among all markers.

Both *DLX3* (F) and *OTOF* (G) were co-expressed with *NMU* in ENs of the adult human ileum. Arrowheads mark instances of colocalization for all markers.

(H) Similar to mice, the human genes *DLX3* and *OTOF* showed strong colocalization in EN, although collectively, there appeared to be a greater proportion of neurons co-expressing these markers in humans than mice. Arrowheads indicate instances of colocalization for all markers.

**Supplemental Figure 7.** Cross-species comparison of myenteric IPAN markers using HCR FISH with intestinal tissue from healthy adult humans and mice.

(A-B) *NMU* is expressed selectively in myenteric IPANs of the adult human ileum (A) and colon (B). Arrows label IPANs expressing *NMU*. Arrowheads mark IPANs that lack *NMU* expression. Asterisks mark *NMU*+ neurons that are not IPANs.

(C-D) *KLHL1* was not observed in IPANs from the ileum (C) and colon (D). Arrowheads mark IPANs that are *KLHL1*[-]. Arrows mark *KLHL1*+ neurons that are not IPANs.

(E) *KCNH7*, a marker co-expressed in *KLHL1*+ neurons (from clusters 6 and 7 in mice), is not expressed in IPANs, but is detected in many neurons in the same field. Arrowheads point to a representative IPAN that does not express *KCNH7*.

(F) In the colon of mice, *Nxph2* is expressed in a subset of *Klhl1*+ neurons devoid of Cdh9 expression (arrowheads), consistent with snRNA-Seq data.

(G) In the colon of mice, *Cdh9* is expressed in a subset of *Klhl1*+ neurons devoid of

*Nxph2* expression (arrowheads), consistent with snRNA-Seq data.

(H-J) Human *KLHL1*+ ENs express the mouse subtype markers, *CDH9* and *NXPH2*. *CDH9* is expressed in a subset of *KLHL1*+ neurons in the duodenum (H) and ileum (I) of adult humans but was observed in many *KLHL1*[-] neurons. In some samples of human small intestine, *NXPH2* was found to be colocalized with *CDH9* (H), in contrast with the discrete marker gene-expression patterns observed in mice.

Similarly, *NXPH2* is expressed in a subset of *KLHL1*+ neurons in the duodenum (H) and ileum (J) as well as in many *KLHL1*[-] neurons (Supplementary Figure 6B). *NXPH2* expression did not coincide with *NMU*+ neurons in the vast majority of cases (J).

For H-J, Arrowheads mark neurons co-expressing *KLHL1* and *CDH9* and/or *NXPH2*. Arrows mark *KLHL1*+ neurons that do not co-express either *CDH9* or *NXPH2*.

**Supplemental Figure 8.** Refined characterization of human *KLHL1*+ EN via HCR FISH indentifies co-localization with the Type III neuron marker *CALB1*.

(A-B) *KLHL1* is colocalized with nearly all observed *CALB1*+ neurons in the duodenum

(A) and ileum (B). *NXPH2* observed to be colocalized with *KLHL1*/*CALB1* in a minority of neurons from these gut regions, despite being commonly expressed in surrounding myenteric neurons and/or glia. Arrowheads mark *KLHL1*+/*CALB1*+ neurons devoid of *NXPH2* expression. Arrows mark a *NXPH2*+ neuron faintly co-expressing *KLHL1* and *CALB1*.

No instances of *CDH9* colocalization with *KLHL1*+/*CALB1*+ neurons in the colon were observed, despite being expressed in many surrounding neurons. Arrowheads mark instances of colocalization between *KLHL1* and *CALB1* that do not express *CDH9*.

**Supplemental Figure 9.** Identification of regionally expressed marker genes for putative subtypes of mouse myenteric neurons.

Mouse EN subtype markers were directly identified independent of matching to human LCM RNASeq data or removing genes expressed in glia or muscle. The top 3 highest-scoring marker genes were derived from mouse snRNA-Seq data and plotted individually as split-out panels for the duodenum (A), ileum (B), and colon (C). A modified scoring approach was used, as described in Supplementary Methods, which enriched for the selection of genes restricted to particular segments.

(D) For ease of visualizing the regional expression of marker genes in each cluster from panels A-C, marker genes were plotted as a heatmap. Each row of the heatmap depicts the expression of each marker gene from the single cluster in which it was expressed most highly. The cluster being plotted is indicated in brackets at the right of each gene symbol. Marker genes are arranged by the order of their regional specificity.

**Supplemental Figure 10.** Cross-species comparison identifies regionally expressed marker genes in myenteric EN subtypes.

(A) Violin plot showing absence of *Cckar* expression in the ileum of mice, with only 1% of neurons in cluster 6 containing significant expression.

(B) *Cckar* was confirmed to be colocalized with *Nxph2* (cluster 6 neurons) in the mouse ileum in all observed cases (arrowheads).

(C) *CCKAR*+ neurons express *OTOF* (arrowheads) in the duodenum, but are not IPANs, because no instances of *CCKAR* expression were observed in *CALB2*+/*OTOF*+ neurons (arrows).

**Supplemental Figure 11.** Regional expression of subtype-specific marker genes mapped across the intestine.

(A) Expression of Cckar in mouse snRNA-Seq clusters, split-out into separate panels by intestinal segment. Numbers label each cluster ID. In the duodenum, strong expression of *Cckar* was observed in the majority neurons of cluster 6 as well as in a relatively large number of neurons in clusters 3 and 0, albeit proportionally much fewer than cluster 6. In contrast, the ileum and colon contained extremely few *Cckar*+ neurons in any cluster.

(B) Expression of *Cckar* in cluster 6 was observed in ~90% of nuclei in the duodenum, but was only observed in ~1% of nuclei in the ileum and colon. Distribution of *Cckar* expression in nuclei of the ileum and colon in UMAP space indicated that *Cckar+* neurons (purple) and *Cckar*[-] neurons (gray) represent the same subtype of neuron and are similar to *Cckar*+ neurons in the duodenum, given uniform distribution and equal intermingling on this cluster.

(C) Cartoon diagram illustrating the concept of regional gene expression within a subtype of EN. The intestine is represented as a pink cylinder and depicts an example of an EN subtype that stably expresses a set of genes throughout the intestine (cluster 6 expresses *Klhl1*, *Kcnh7*, *Nxph2*, and *Nefl*, etc.). The chart above the intestine displays the percentage of neurons from cluster 6 that were observed in different regions of the intestine. In the duodenum, Cckar is expressed in the majority of neurons of cluster 6, with extremely few *Cckar+*/*Nefl+*/*Klhl1+* neurons observed in the ileum and colon. Note that the gradual reduction in the percentage of neurons from the proximal duodenum, through the jejunum and to the distal ileum is arbitrary. No quantification was performed *in vivo* or in different subregions of each gut segment.

## References

Author names in bold designate shared co-first authorship.

