## Supplementary Figure Compilation for "Combinatorial transcriptional profiling of mouse and human enteric neurons identifies shared and disparate subtypes *in situ*"

#### Supplemental Figure 1

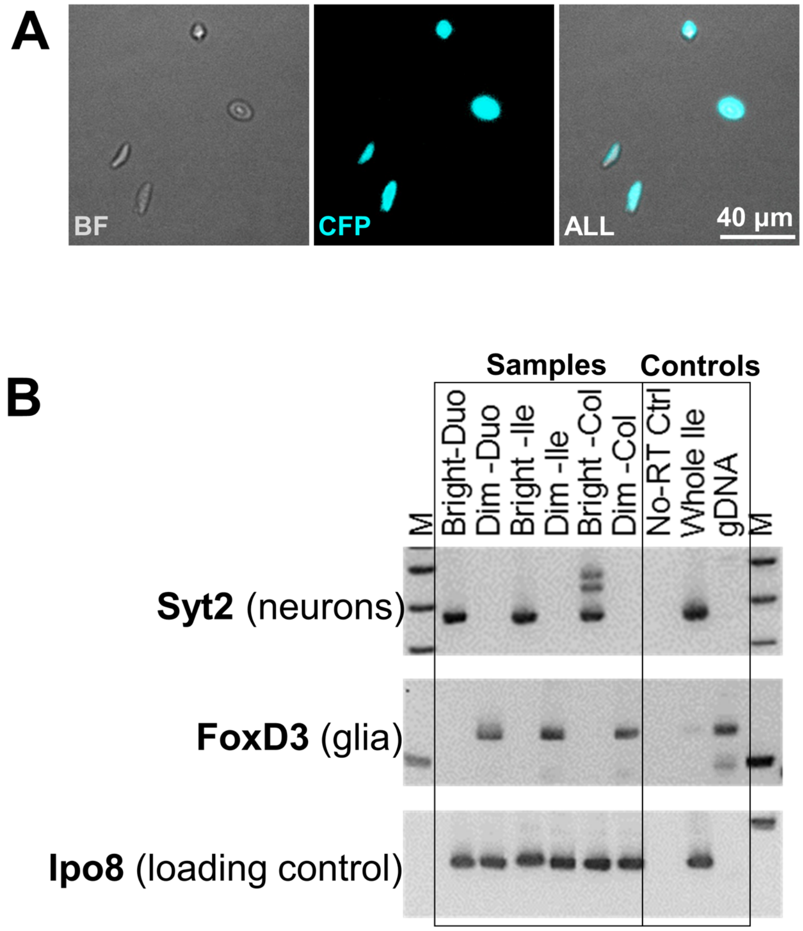

### Supplemental Figure 2

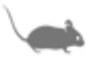

| Sn-RNA-Seq run summaries | Duodenum | Ileum | Colon | All |
|---|---|---|---|---|
| Approx. Nuclei Encapsulated | 12884 | 24413 | 17599 | 54896 |
| 10X Total Runs | 2 | 4 | 3 | 9 |
| inDrop Total Runs | 2 | 2 | 2 | 6 |
| Total Runs | 4 | 6 | 5 | 15 |
| 10x Total Nuclei | 5394 | 8291 | 14271 | 27956 |
| inDrop Total Nuclei | 1756 | 3570 | 3328 | 8654 |
| Total Nuclei Pre-cleaning | 7150 | 11861 | 17599 | 36610 |
| Total Nuclei Post-cleaning * | 6217 | 8379 | 10612 | 25208 |
| Total Reads 10x | 7.8E+08 | 1.8E+09 | 9.5E+08 | 3.6E+09 |
| Total Reads inDrop | 1.7E+08 | 1.3E+08 | 2.5E+08 | 5.5E+08 |
| Total Reads | 9.4E+08 | 2.0E+09 | 1.2E+09 | 4.1E+09 |

### Supplemental Figure 3

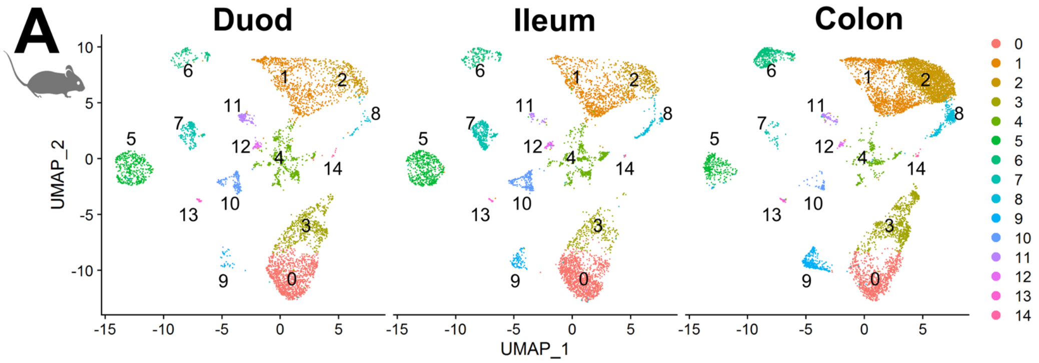


#### B Gut segment compostion for each cluster (%)

| Cluster | 0 | 1 | 2 | 3 | 4 | 5 | 6 | 7 | 8 | 9 | 10 | 11 | 12 | 13 | 14 |
|---|---|---|---|---|---|---|---|---|---|---|---|---|---|---|---|
| Duo | 24.2 | 17 | 7.3 | 13 | 11.6 | 9.9 | 2.4 | 4.4 | 0.9 | 0.7 | 3.8 | 2.9 | 1 | 0.6 | 0.2 |
| Ile | 23.9 | 17 | 7.3 | 6.1 | 11.1 | 10.4 | 3.4 | 8.2 | 3.8 | 1.8 | 5 | 0.9 | 0.9 | 0.3 | 0.1 |
| Col | 11.8 | 18.2 | 30.3 | 11.6 | 3.9 | 4.1 | 6.4 | 0.6 | 4.9 | 5.3 | 0.6 | 0.9 | 0.6 | 0.5 | 0.4 |

#### C Putative subtype identities and top cluster markers

| Cluster | Known Markers | Proposed Subtype Identity | Top 3 Mouse | Top 3 Human-Match |
|---|---|---|---|---|
| 0 | Calb2/Chat/ Tac1+/- | Excitatory longitudinal muscle motor neurons | Brinp2/Fbxw15/Specc1 | Oprk1/Brinp2/Tmem132c |
| 1 | Nos1/Vip | Inhibitory longitudinal muscle motor neurons | Ass1/Cygb/Col25a1 | Ngb/Gsg1l/Col25a1 |
| 2 | Nos1/Vip/Npy[+/-] | Inhibitory circular muscle motor neurons | Gm4876/Gm16083/Dsc2 | Vwa5b1/Sv2b/Oprd1 |
| 3 | Chat/Tac1 | Excitatory circular muscle motor neurons | Tac1/Necab2/Htr2b | Penk/Sncb/Olfm3 |
| 4 | - | Unassigned | Slc24a3/Myh11/Carmn | Cemip/Kcnip1 |
| 5 | Calcb/Calb2[+/-] | IPAN | Nmu/Pcdh10/Cysltr2 | Nmu/Pcdh10/Cbln2 |
| 6 | Nefl/Calb1[sparse] | IPAN | Nxph2/Cckar/Eif3h | Skap1/Nefl/Nxph2 |
| 7 | Nefl | Viscerofugal? (Type I/'simple') | Cdh9/Zim1/B230209E15Rik | Cdh9/Mgat4c/Klhl1 |
| 8 | Nos1/Gal/Sst(low) | Unassigned | *No selective markers | *No selective markers |
| 9 | Sst/Calb2/Th | Descending interneurons (Filamentous) | Adamts1/Gm30382/Irf1 | Sst/Galnt5/Pantr1 |
| 10 | Chat/Nos1/Gad2 | Descending interneurons (Type I) | Neurod6/AI593442/Bcr | Trhde/Bean1/Neurod6 |
| 11 | - | Unassigned | C3/Igfbp6/Upk3b | Myrf/Nkain4/Cldn15 |
| 12 | - | Unassigned | Cdh19/Apoe/Col11a1 | Col11a1/Tmprss5/Car12 |
| 13 | - | Unassigned | Reg3b/Pigr/Epcam | Pigr/Klf5/Cdh1 |
| 14 | - | Unassigned | Ccl21a/Mmrn1/Pecam1 | Klhl4/Arap3/Radil |

#### D Proposed cluster subtype identities

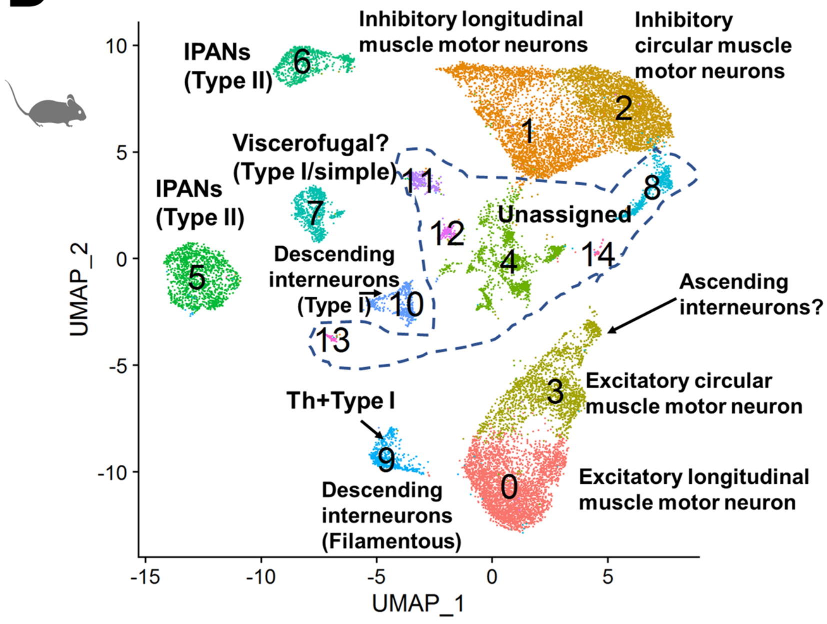

### Supplemental Figure 4

**A**

| Cluster | KEGG Function | q_value |
|---|---|---|
| 4 | Oxytocin signaling pathway | 3.8E-08 |
| 4 | Axon guidance | 6.0E-03 |
| 4 | Dopaminergic synapse | 4.8E-02 |
| 8 | Synaptic vesicle cycle | 6.5E-02 |
| 8 | Glutamatergic synapse | 1.3E-01 |
| 8 | Long-term potentiation | 1.6E-01 |
| 11 | Axon guidance | 1.0E-02 |
| 12 | Dopaminergic synapse | 8.6E-02 |
| 12 | Glutamatergic synapse | 1.0E-01 |
| 12 | Axon guidance | 1.3E-01 |
| 13 | Alzheimer disease | 1.2E-08 |
| 13 | Huntington disease | 1.2E-08 |
| 13 | Serotonergic synapse | 2.9E-01 |
| 14 | Axon guidance | 1.8E-03 |
| 14 | Oxytocin signaling pathway | 2.9E-01 |
| 14 | Long-term depression | 5.3E-01 |

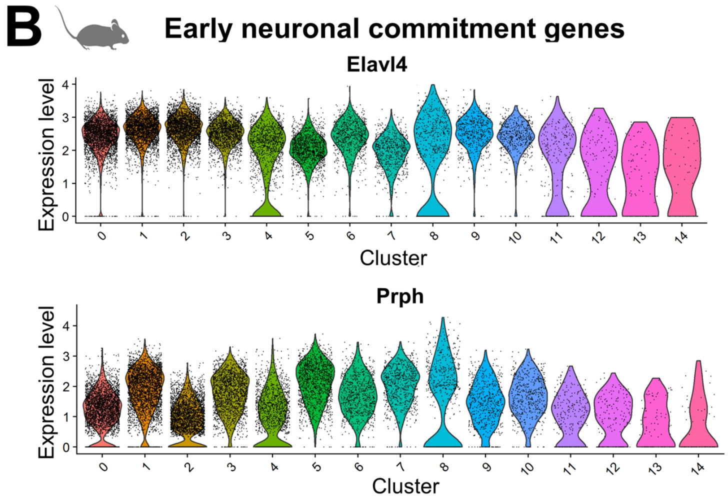


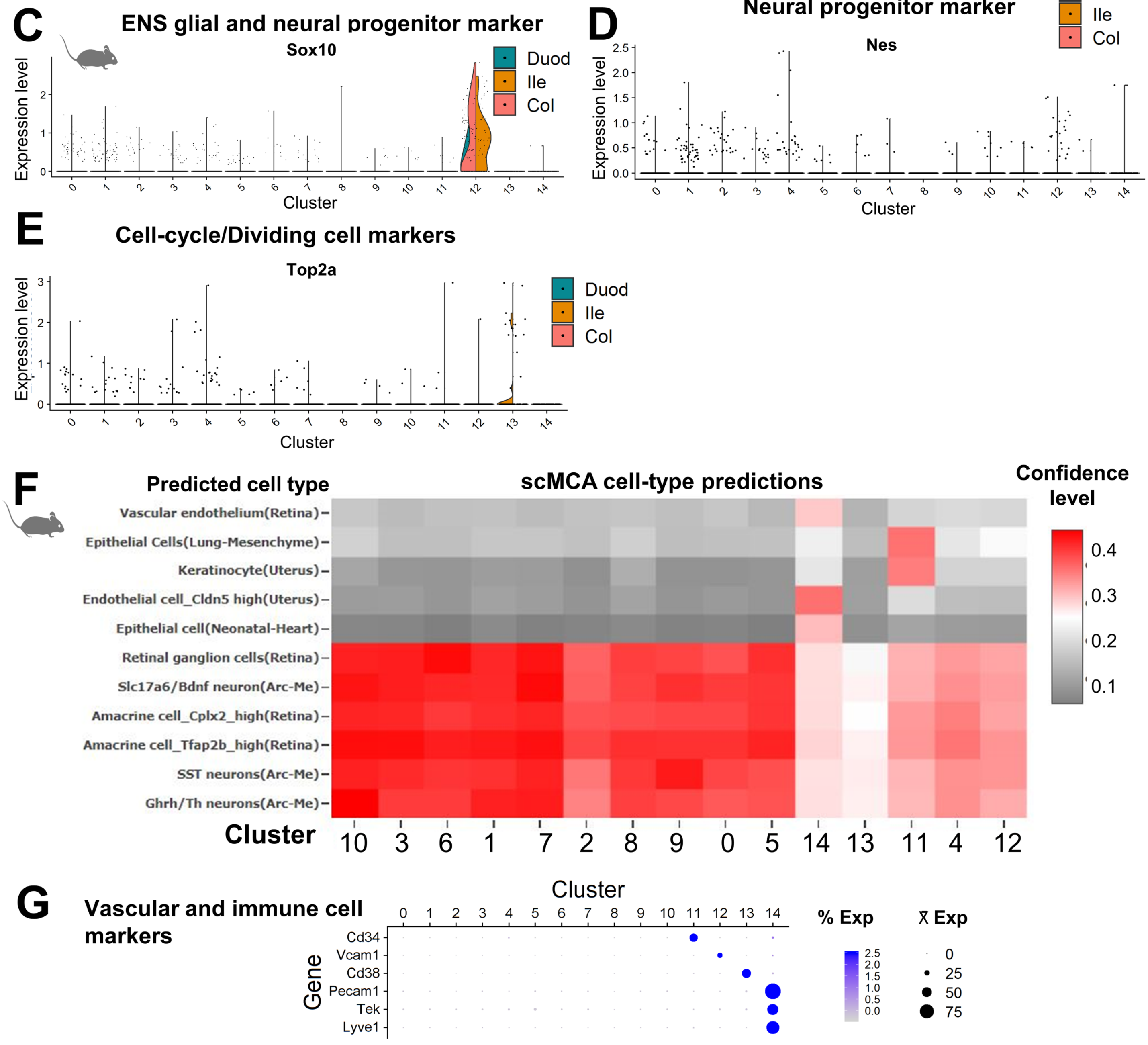

### Supplemental Figure 5

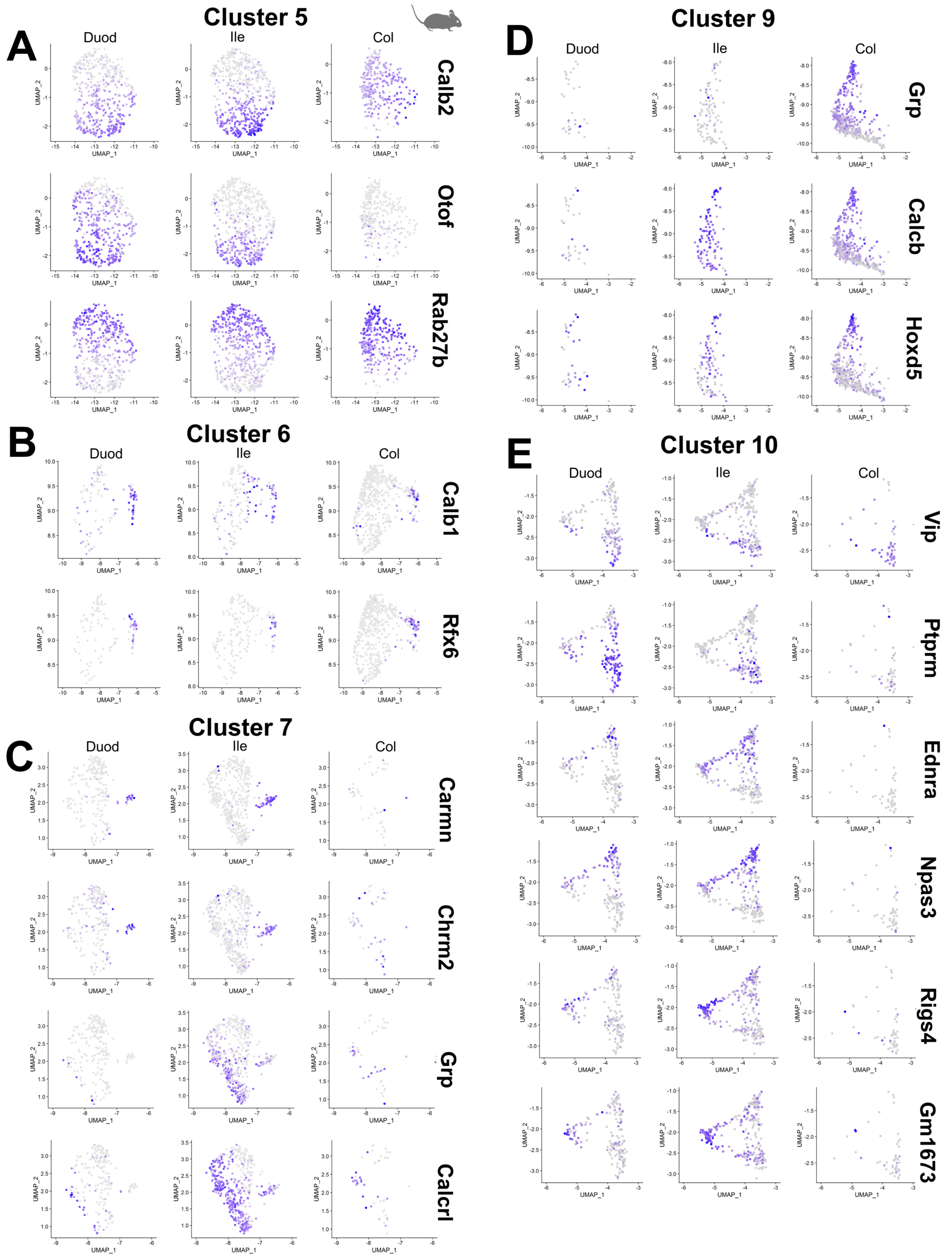

### Supplemental Figure 6

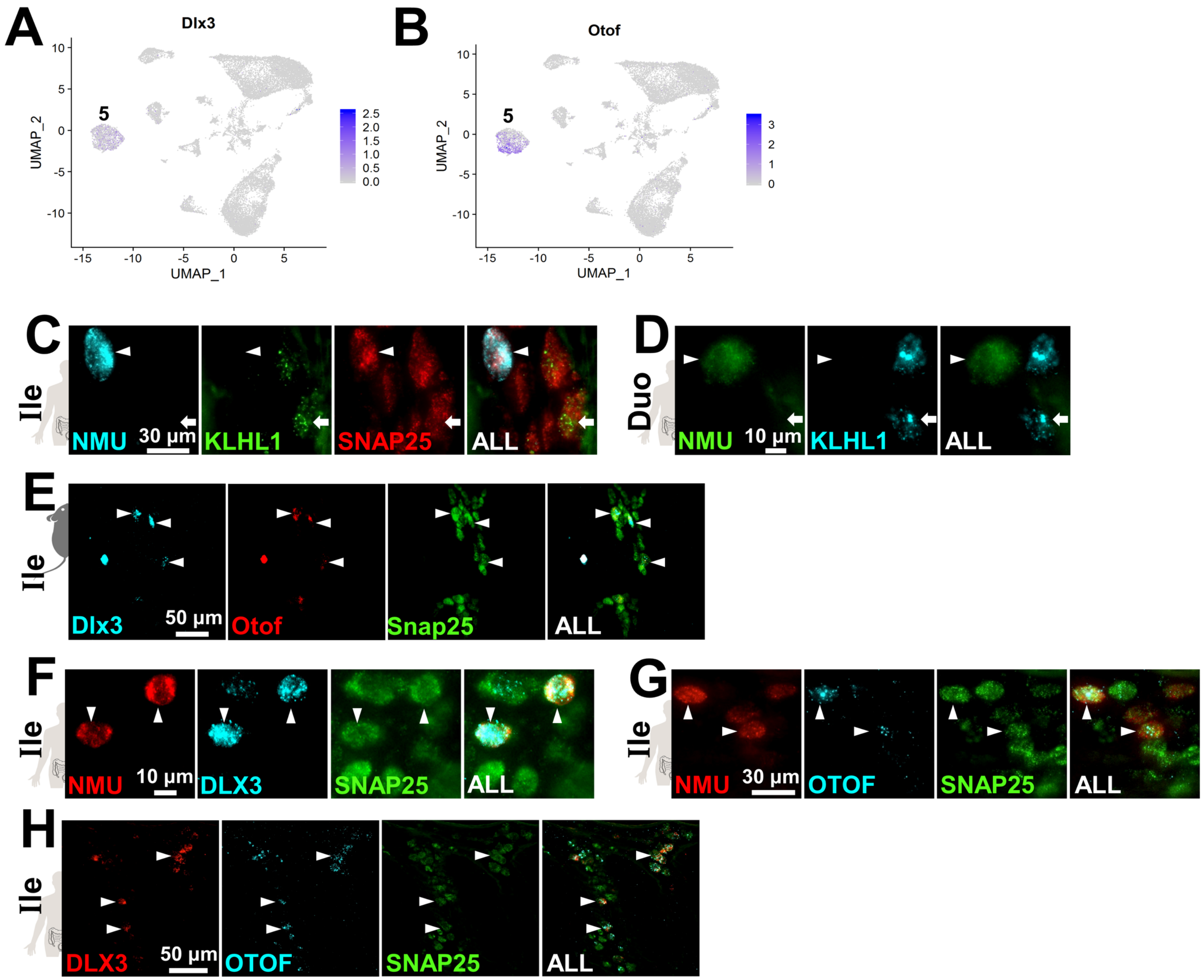

### Supplemental Figure 7

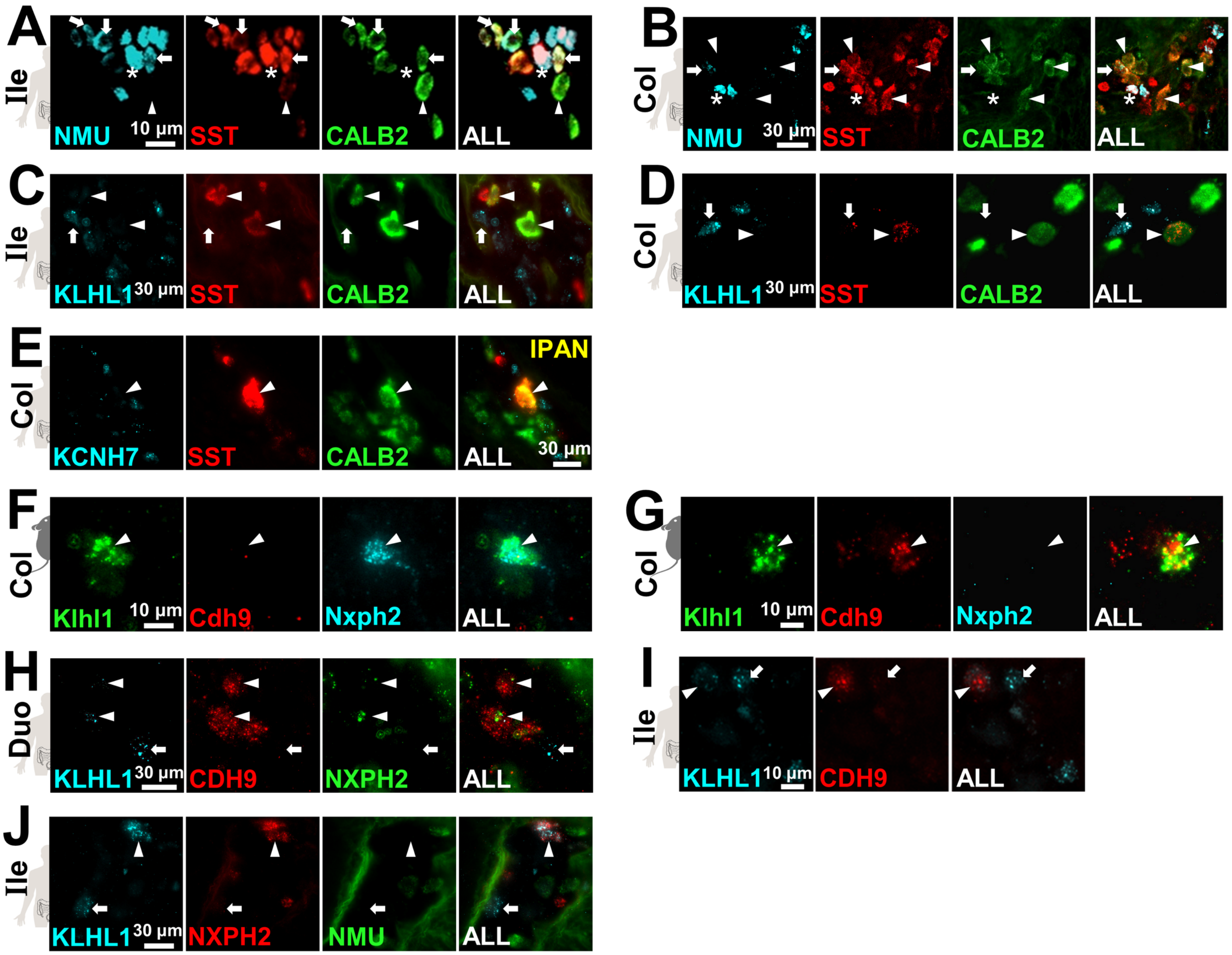

### Supplemental Figure 8

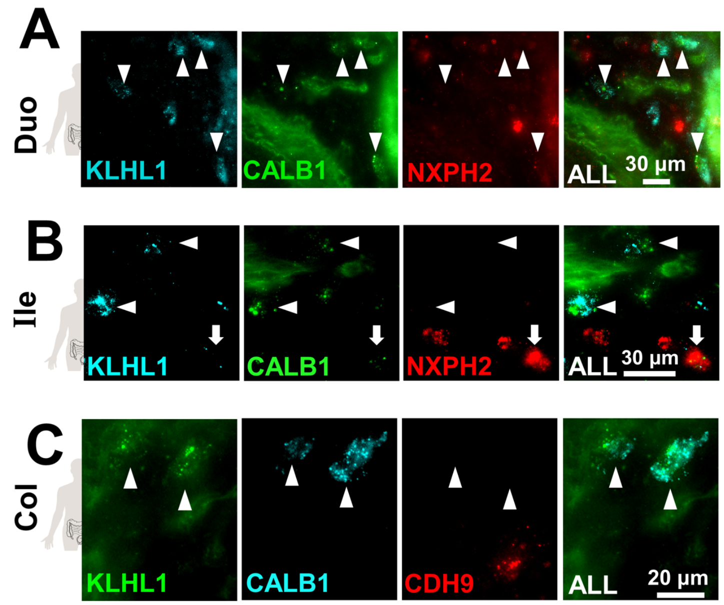

#### Supplemental Figure 9

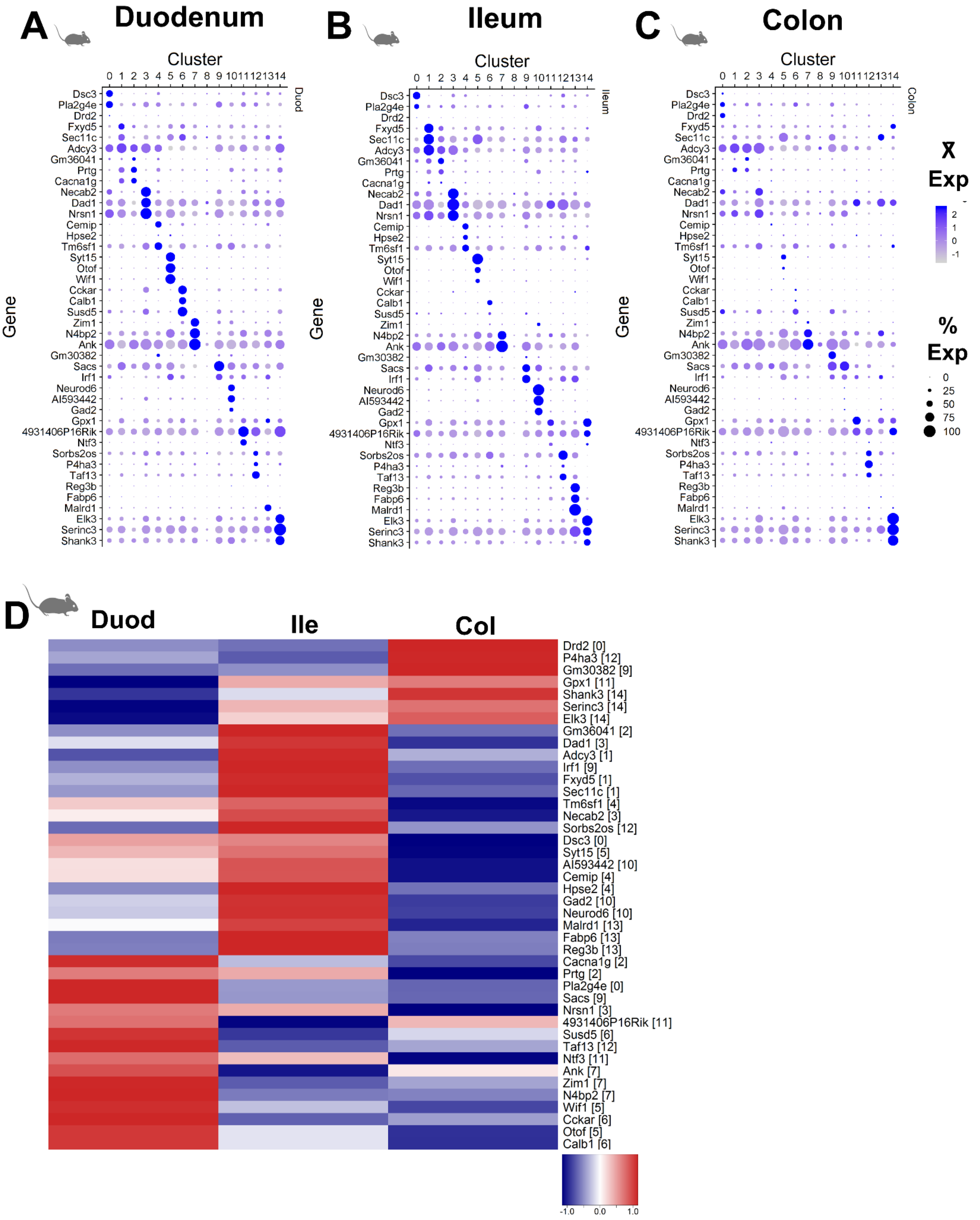

### Supplemental Figure 10

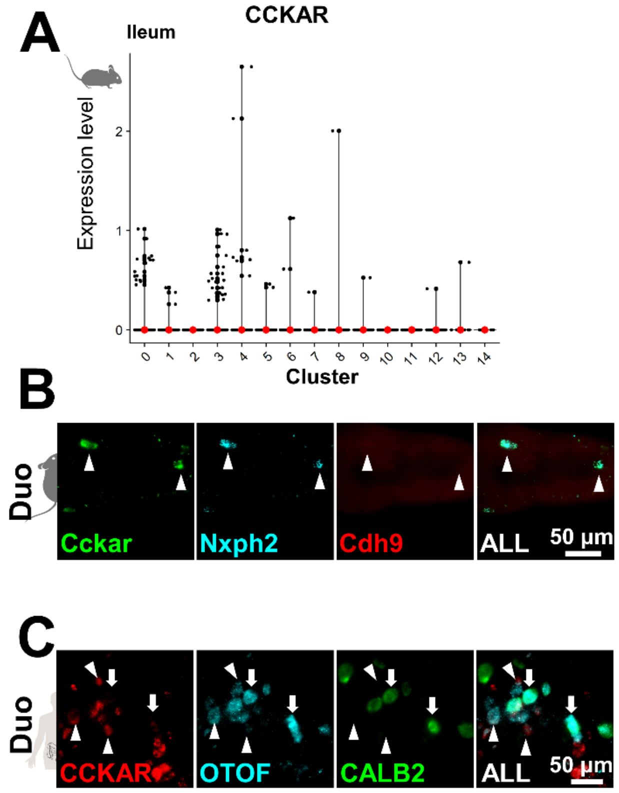

### Supplemental Figure 11

**A** All clusters

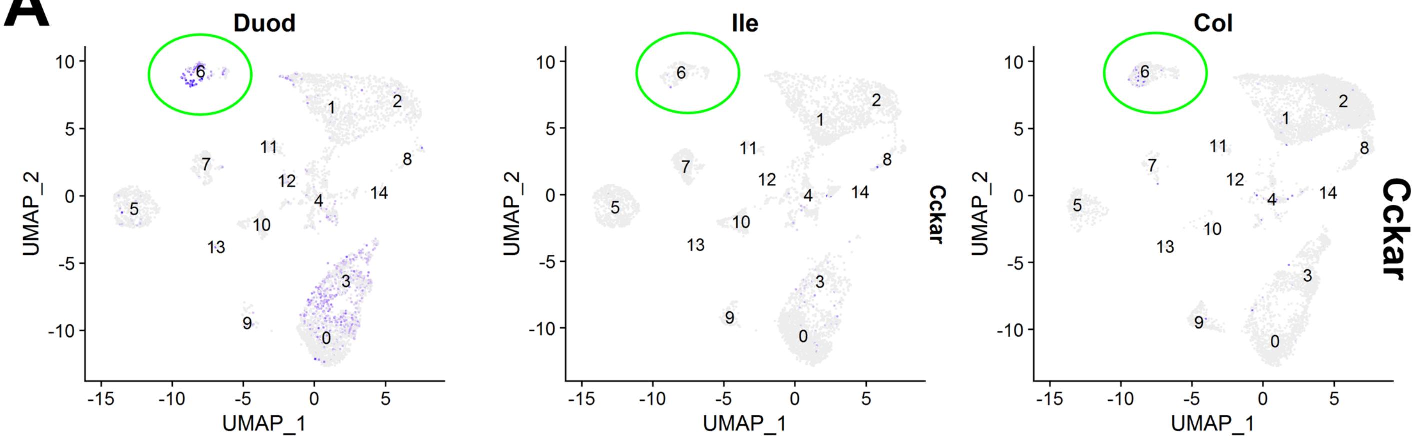


**B** Cluster 6 only

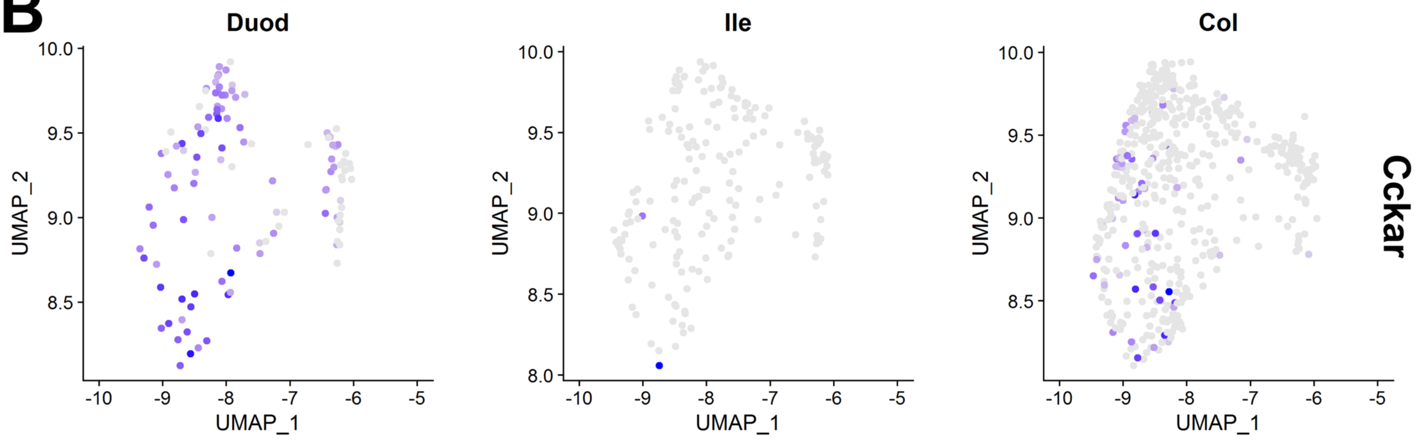


**C** Regional expression of Cckar in neuronal cluster 6 in mice

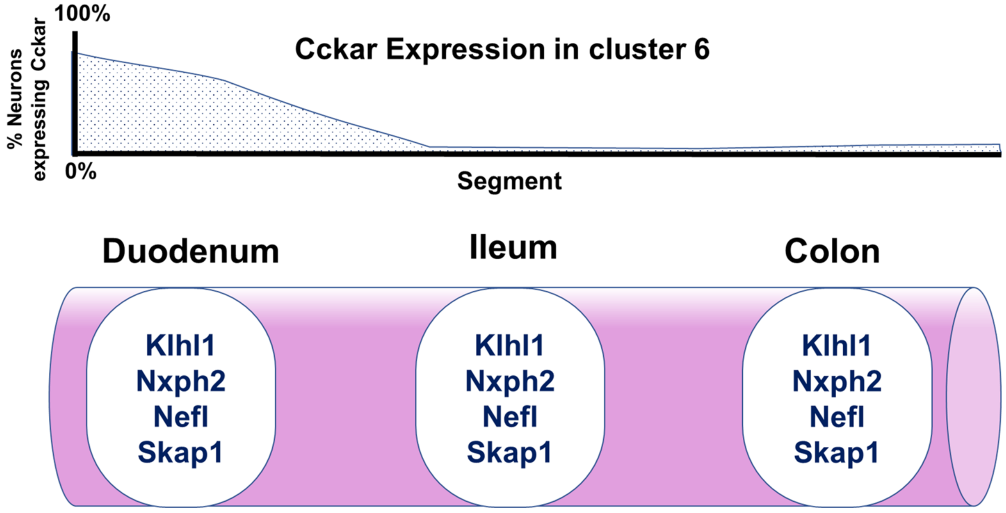
