## Supplementary Methods for "Combinatorial transcriptional profiling of mouse and human enteric neurons identifies shared and disparate subtypes *in situ*"

**Nuclei isolation**

Minced tissue (3x5-mm segments) from the muscularis externa of mouse duodenum, ileum, and colon was homogenized while submerged in 2-mL of nuclei lysis buffer consisting of Nuclei EZ lysis buffer (Sigma), 0.2 U/μL RNAse inhibitor (Roche, Protector®), and 1x Protease inhibitor cocktail, EDTA-free (Roche complete mini). The lysate was filtered through a 40-μm nylon strainer into a 5-mL round-bottom collection tube leaving all un-minced tissue in the Dounce mortar. The filter was rinsed repeatedly with 1x RNAse-free DPBS with 2mM Mg2+. All filtrates were centrifuged at 500 x g at 4^o^C, forming a compact, but loose pellet. Supernatant was completely removed before adding 100μL of ice-cold RNAse-free 1x DPBS (with 2 mM Mg2+) to each nuclei pellet. Pellets were resuspended using a normal-bore pipette tip just until any large aggregates were dispersed. Samples were then either fixed with DSP (Lomant’s reagent) or were processed immediately with FACS. Nuclei were collected into Eppendorf LoBind protein plates pre-coated with 1% BSA in 1X PBS.

**RT-PCR and Preamplification: Isolated nuclei retain RNA signatures of cellular identity**

To confirm whether FACS-purified nuclei from bright and dim populations exhibited transcripts representative of neurons and glia, respectively, we performed RT-PCR. cDNA was pre-amplified for 14 cycles (TaqMan^TM^ PreAmp Master Mix, Applied Biosystems). Subsequently standard PCR reactions were performed (94^o^C x 5 min followed by 36 cycles of 94 ^o^C x 30 s, 52.4 ^o^C (or 55 ^o^C for TaqMan probes) x 30 s, 72^o^C x 30 s, and ending with 72 ^o^C x 10 min using AmpliTaq® DNA Polymerase (Applied Biosystems). The presence or absence of PCR products was visually assessed on 10% acrylamide gel (Fig S2A). Controls included: no template, RNA subjected to processing without addition of reverse transcriptase (no RT), and total genomic DNA

**Reversible fixation with DSP for bulk nuclei and snRNA-Seq**

Fixation of nuclei with DSP was performed using slight modifications to a published method^1^. Within five minutes of resuspending nuclei, DSP was solubilized in 100% anhydrous DMSO (Sigma) at a final concentration of 25 mg/mL to form a “25X” stock. The 25X stock was then quickly added dropwise to DPBS (with 2 mM Mg2+) inside of a 50-mL conical tube being vortexed continuously at medium speed. After a few brief centrifugations to thoroughly mix the solution, the 1x DSP solution was then immediately filtered through a 0.2-μm PES membrane syringe filter and collected into a 15-mL conical tube. If any precipitates were visible prior to filtering, the solution was remade, because this consistently resulted in crystal formation in the filtrate. This solution was observed to be stable at room temperature for 3-5 minutes. As soon as nuclei were resuspended, 1.25 mL of 1x DSP was added to each chilled nuclei pellet to achieve a final DSP concentration of ~0.93 mg/mL. Samples were gently inverted and then incubated for 30 minutes at RT (26°C) on a rotating platform to prevent nuclei aggregation. To quench residual DSP, Tris-HCl was then added at a final concentration of 20 mM. Samples were kept at 4°C to avoid the precipitation of DSP on ice prior to FACS. Of the 16 nuclei samples that were successfully sequenced, 10 were fixed and five were left unfixed. Unfixed samples included inDrop Run 3445 (all segments) and 10x Run 0052 (one ileum and one colon sample).

**inDrop nuclei encapsulation and sequencing (additional information)**

Syringes and tubing were blocked with 1% BSA (New England Biosciences, #B9000S) prior to loading. The tubing was loaded first with glial nuclei (“Dim” sorted population) to fill the dead volume, then neuronal nuclei (“Bright” sorted population). inDrop utilizes CEL-Seq in preparation for sequencing^2^ and is summarized as follows: 1) Reverse transcription (RT) (supplemented with 30µM DTT), 2) ExoI nuclease digestion (no HinFI), 3) SPRI purification (SPRIP), 4) Second strand synthesis, 5) T7 in vitro transcription linear amplification, 6) SPRIP, 7) short RNA fragmentation, 8) SPRIP, 9) Primer annealing, 10) RT, 11) SPRIP and 12) library indexing PCR (doubled reaction size), 13) SPRIP. Number of nuclei encapsulated was calculated by observing the single nuclei suspension loading rate multiplied by bead loading efficiency during the duration of encapsulation. The largest sample contained an estimated 5000 nuclei, while the smallest contained an estimated 100 nuclei. Following library preparation, the samples were sequenced using a Nextseq 500 (Illumina) with a 150bp paired-end sequencing kit in a customized sequencing run, as described in Methods.

**Single-Nucleus RNA-Seq data processing for raw matrices**

The number of usable nuclei sequenced from each run was determined by plotting the cumulative summed UMIs and identifying the radius of least curvature, representing the estimated point at which droplets transition from solely containing ambient RNA to containing encapsulated, intact nuclei ^3^. Only droplets containing singlets were included in the final analysis, as determined using the DoubletFinder R-package^4^. Runs from the duodenum, ileum, and colon were separately aggregated and batch-corrected using Seurat’s integration method with default settings^5^. Next, Louvain clustering and Uniform Manifold Approximation and Projection (UMAP) analysis was performed within Seurat. After identifying clusters, marker gene lists were generated using differential expression analysis with the Wilcoxon rank-sum test and Bonferroni p-value correction (q ≤0.05).

**Bulk RNA-Seq data analysis for human enteric ganglia and nuclei from mouse ENs**

Base calls and demultiplexing were performed with Illumina’s bcl2fastq software with a maximum of one mismatch in the indexing read. RNA-Seq reads were then aligned to their respective Ensembl release 76 human or mouse top-level assemblies with STAR version 2.0.4b. Gene counts were derived from the number of uniquely aligned, unambiguous reads by Subread:featureCount version 1.4.5. Isoform expression of known Ensembl transcripts were estimated with Sailfish version 0.6.13. Sequencing performance was then assessed for the total number of aligned reads, total number of uniquely aligned reads, and features detected. The ribosomal fraction, known junction saturation, and read distribution over known gene models were quantified with RSeQC version 2.3.

All gene counts were then imported into the R/Bioconductor package edgeR and TMM normalization size factors were calculated to adjust for differences in library size across samples. Ribosomal genes and genes not expressed in the smallest group size were excluded from further analysis. The TMM size factors and the matrix of counts were then imported into the R/Bioconductor package Limma and weighted likelihoods based on the observed mean-variance relationship of every gene and sample were then calculated for all samples with the voomWithQualityWeights function. The performance of all genes was assessed with plots of the residual standard deviation of every gene to their average log-count with a robustly fitted trend line of the residuals. Differential expression analysis was then performed to analyze for differences between enteric ganglia and intestinal muscle between all gut segments. Results were filtered to select genes with Benjamini-Hochberg false-discovery rate adjusted p-values (q) of q ≤ 0.05. The R/Bioconductor package heatmap3 was used to display heatmaps across groups of samples.

For mouse neuronal nuclei samples, two samples of ENs and glia were sequenced for the duodenum, ileum, and colon, with the exception of colon only having a single glial sample.

**SnRNA-Seq: Dimensionality reduction and clustering of individual gut segments**

SnRNA-Seq count matrices were processed using the Seurat package (version 3.1.2, R 3.6.2 library). Separate Seurat objects were generated for each intestinal segment using nuclei collected from the duodenum, ileum, or colon. Only one read per cell was needed for a gene to be counted as expressed. After merging raw matrices together, resulting gene expression matrix was scaled, normalized to 10,000 molecules per cell and log-transformed according to Macosko et al^6^. The top 2000 highly variable genes were used for principal component analysis (PCA). Batch correction was performed using Harmony^7^, because Seurat’s integration approach^5^ did not fully correct for batch differences of cell types known to have similar morphology, neurophysiology, and marker genes between segments, such as *Nos1*+ neurons (data not shown). In total, 15 clusters were identified, representing at least 15 major discrete neuronal subtypes across the entire intestine.

We next inspected whether any clusters had uneven distributions of cells across batches that may have aberrantly resulted from batch correction or other sources of error during the sequencing and/or analysis process. Sampling of neuronal subtypes from the intestine of Phox2b-CFP mice was expected to be relatively consistent for all batches. The variance for the expected nuclei per cluster from each batch was estimated using 99% confidence intervals derived from nuclei counts in each snRNA-Seq cluster. Clusters were flagged if nuclei counts fell outside of the confidence interval. Clusters were removed from subsequent analysis if the majority of the samples (>50%) were flagged in the single cluster. When performing Harmony, all batch-corrected samples met the above criteria and had a relatively even representation of nuclei from the duodenum, ileum, and colon and no clusters were excluded. This was not the case for the other batch-correction approaches, such as Seurat integration^5^, in which several nuclei clusters were flagged as not meeting the expected proportions of nuclei. Flagged clusters were found to likely consist of neuronal progenitors. Given the uncertainty of batch-correction procedures when examining unknown subtypes of neurons, we examined all putative progenitor clusters with the ClusterProfiler R package, to ensure that these subtypes are biologically relevant and observed that all clusters had significant KEGG terms and scMCA identity predictions relating to the nervous system (Supplemental Figure 5A,F)

To ensure that all clusters generated after batch correction with Harmony represent biologically-meaningful subtypes, we first removed doublets using the DoubletDecon R package^8^. Expected doublet rates were ~5%^9^, so we tested doublet corrections at varying stringency levels that removed 5-20% of all nuclei in the dataset as putative doublets and examined whether any clusters consisted mostly of doublets. However, the putative doublets were distributed relatively evenly across all clusters, indicating there were no artifactual clusters generated from the presence of doublets. We applied the most stringent doublet-removal settings when moving forward to marker gene identification because no clusters were lost during the doublet-removal process and each cluster had strong representation from each gut segment.

We further examined all putative progenitor clusters with the ClusterProfiler R package, to ensure that these subtypes are biologically relevant and observed that all clusters had significant KEGG terms and scMCA identity predictions relating to the nervous system (Supplemental Figure 4A,F).

To identify all neuronal subtypes present across the entire intestinal tract, we performed dimensionality reduction and unsupervised clustering using the UMAP algorithm^10^ using the first 50 principal components. All clustering was unsupervised, without the use of driver genes. Local network graphs were calculated using the shared nearest neighbors (SNN) method and clusters were assigned using the Louvain clustering algorithm^5^. UMAP plots visualized the single cells in two-dimensional space based on expression signatures of the variable genes. Over/under clustering was verified using gene expression heatmaps and with visual inspection of UMAP plots to examine expression of known subtype markers within discrete clusters. The resolution parameter was adjusted between 0.3 to 0.6 and data were re-clustered until snRNA-Seq data had distinct clusters for at least all known published markers.

**Marker gene identification and prioritization**

A bioinformatics screening procedure was used to select the optimal marker genes for discrete neuronal subtypes. Differential expression was performed separately for all three gut segments using the same embeddings and clustering from the Harmony-corrected, merged dataset described above. First, significant marker genes were identified for each cluster using the Wilcoxon rank-sum t-test for each cluster using a “one-vs-all” cluster comparison. Gene lists from each cluster represent markers for putative neuronal subtypes. Markers were filtered to include only genes with a Bonferroni-adjusted p-value of ≤ 0.05. Human gene orthologs were identified for the respective mouse genes using the MouseMine homolog-matching tool (mousemine.org). Human orthologs were then matched against the list of genes that were significantly enriched in enteric ganglia relative to intestinal muscle. Genes with conserved expression between mice and humans were selected for further investigation. We next sought to identify markers that would perform optimally with *in situ* hybridization. This involved identifying markers expressed specifically in neurons, with minimal expression in enteric glia and muscle. Neuronal subtype marker genes were discarded if expression was observed in single-cell data sets of mouse enteric glia^11^ (>5% of cells) or intestinal muscle (>10% of cells in a cluster). Next, marker genes were filtered to select and retain genes with a log2-fold change of ≥ 3. Finally, marker genes were ranked by their suitability as a marker of both human and mouse neuron subtypes using a weighted algorithm. The highest score was given to markers that were 1) expressed only in a single cluster, 2) highly expressed, 3) expressed in a high percentage of nuclei within a given cluster, 4) expressed minimally in all other nuclei clusters, 5) expressed highly in human ganglia, 6) minimally expressed in human intestinal muscle.

Markers were then unified based on their segment of origin and were assessed for their specificity for a single cluster in mice or expression in a particular gut region in both mice and humans. Most genes detected were expressed in all three segments, but many were identified with regional expression, as described below (Figure 7,Supplementary Figures 9 and 11).

A caveat with this approach is that the comparison of mouse neuron subtypes with human ganglia is not a direct neuron-to-neuron comparison. Differences in expression within non-neuronal cells in enteric ganglia have the potential to overpower true similarities between neurons across species. Conversely, many genes that appeared to work well based on the expression within whole ganglia were inconsistent. Promising cluster-specific markers that were not reliably detected in neurons using HCR included: NPY5R, CCK, RFX6, WIF1 and VWCL2 (data not shown). While it is possible HCR probes were simply nonfunctional for these genes, this is highly unlikely given the number of occurrences and more likely reflects differences in expression between species.

**Segment-specific marker gene identification and prioritization**

Further screening was performed to identify markers that are specific to neurons of individual gut segments. First, markers derived in the above process were passed through an intra-cluster differential expression test to identify cluster-specific differences in gene expression. This was achieved by performing the Wilcoxon rank-sum test with Bonferroni correction for each cluster, separately and comparing one gut segment vs. all (i.e. colon vs. duodenum + ileum). All genes with a significant difference (p-adj ≤ 0.05) at this level were flagged and labeled with relevant indication: i) colon-specific, small-intestine-specific (enriched in either the duodenum or ileum) or present in all segments (having no significant difference in expression between any gut segments). This process was repeated for each cluster across the entire dataset. Only genes specific to the small intestine or colon were carried forward for segment-specific analysis (Supplementary Figure 9). Marker genes were prioritized in a manner similar to the above approach. Markers were ranked using the previously-calculated scores taken from only the segment with the highest level of expression in the specified cluster.

We visualized several genes appeared to be well-conserved segment-specific markers for human ENs based on snRNA-Seq data, including: WIF1, CCK, and CCKAR. Although probes designed to target mouse orthologs for these genes were consistent with snRNA-Seq data, the corresponding human probe set did not work as expected for WIF1 and CCK. Oddly, probes to these genes yielded high background signal in ganglia, with little-to-no additional expression in neurons. For this reason, we moved forward focusing on CCKAR for subsequent studies.

**Collection and preparation of mouse muscularis externa tissue sections**

Intestinal tissue was collected from adult mice (age 6-7 weeks) and the muscularis externa was peeled away from the submucosa in chilled 1X DPBS (with Mg^2+^/Ca^2+^). Isolated gut muscle strips of duodenum, ileum, or colon were placed into a cryomold pre-filled with Tissue Freezing Medium™ (TFM) after cutting longitudinally down the length of the intestinal tube to generate a flat sheet of tissue. These rectangular strips were submerged in TFM and flash-frozen atop a slurry of 2-methylbutane and crushed dry ice. 10-μm cryosections were collected onto standard charged glass slides after allowing slides to quickly cool inside the -20^o^C cryostat chamber before adhering the tissue sections. Samples were then fixed and dehydrated as described for human tissue below.

**Collection of human intestinal tissue and preparation of cryosections from the myenteric plexus**

Intestinal tissue was collected from a total of 17 disease-free, adult human organ donors (age 18-35) and prepared as described previously^12^. After flash-freezing tissue, specimens were stored at -80^o^C until use. Cryosections of myenteric plexus were prepared from fresh-frozen intestinal tissue of humans (18-25 μm). After adhering sections to briefly-chilled charged glass slides, the slides were immediately immersed in ice-cold 4% paraformaldehyde (PFA) with 3% sucrose for 30-60 min and subsequently fixed at room temperature for an additional 15 min. Slides were then rinsed in RNAse-free 1X PBS (3x5 mins) and dehydrated in ethanol series. Slides were then stored in anhydrous 100% ethanol at -80^o^ C for later processing.

**Quenching lipofuscin autofluorescence following HCR**

To block autofluorescence of lipofuscin, a natural cellular waste product in lysosomes that complicates imaging of neurons in adult tissues^13^, TrueBlack® dye (Biotium #23007) was applied to tissues after HCR processing using slight modifications from the manufacturer’s protocol. A working solution of TrueBlack® was prepared for each sample using 150 μL of 70% ethanol per sample and 2.75 μL of TrueBlack® stock solution. After applying the stain to each slide for 30s with continuous gentle agitation, the stain was poured off and the slide was de-stained in a 30-mL tube of 45% ethanol with gentle agitation. This step was crucial in retrieving the full intensity of fluorescence for lowly-expressed gene targets. Slides were then rinsed with PBS 3x5 mins before mounting in PBS and thoroughly sealing the coverslips, because the combination of HCR and TrueBlack® was found to be chemically incompatible with standard mounting media. Slides were stored long-term in a humidified chamber at 4^o^ C.
